# A maternal high-fat diet predisposes to infant lung disease via increased neutrophil-mediated IL-6 trans-signaling

**DOI:** 10.1101/2024.03.13.584927

**Authors:** Bodie Curren, Tufael Ahmed, Ridwan B. Rashid, Ismail Sebina, Md. Al Amin Sikder, Daniel R. Howard, Mariah Alorro, Md. Ashik Ullah, Alec Bissell, Muhammed Mahfuzur Rahman, Michael A. Pearen, Grant A. Ramm, Antiopi Varelias, Stefan Rose-John, Robert Hoelzle, Páraic Ó Cuív, Kirsten M. Spann, Paul G. Dennis, Simon Phipps

**Affiliations:** QIMR Berghofer Medical Research Institute, Herston, Queensland, 4006, Australia; School of Biomedical Sciences, Faculty of Medicine, The University of Queensland, Queensland, 4072, Australia; School of Biomedical Sciences, Faculty of Health, Queensland University of Technology, Queensland, 4000, Australia; Christian-Albrechts-Universität zu Kiel, Medical Faculty, Olshausenstraße 40, D-24098 Kiel, Germany; School of Environment, The University of Queensland, Queensland 4072, Australia; Australian Infectious Diseases Research Centre, The University of Queensland, Queensland, 4072, Australia; Centre for Immunology and Infection Control, School of Biomedical Sciences, Faculty of Health, Queensland University of Technology, Queensland, 4000, Australia; School of Biomedical Sciences, Queensland University of Technology, Translational Research Institute, Woolloongabba, QLD 4102, Australia

**Author notes:** Correspondence: Dr. Simon Phipps, QIMR Berghofer Medical Research Institute, Herston, Queensland 4006, Australia. Joint First Authors (Bodie Curren, Tufael Ahmed).

**Keywords:** RSV, bronchiolitis, ALRI, asthma, microbiome, IL-6 trans-signaling, neutrophils, maternal diet, liver

## Abstract

Poor maternal diet during pregnancy predisposes to severe lower respiratory tract infections (sLRI) in infancy, which in turn, increases childhood asthma risk, however the underlying mechanisms remain poorly understood. Here, we show that the offspring of high fat diet (HFD)-fed mothers (‘HFD-reared pups’) developed a sLRI following pneumovirus inoculation in early-life and subsequent asthma in later-life upon allergen exposure. Prior to infection, HFD-reared pups developed microbial dysbiosis and low-grade systemic inflammation (LGSI), characterized by hyper-granulopoiesis in the liver and elevated inflammatory cytokine expression, most notably IL-17A, IL-6 and sIL-6R (indicative of IL-6 trans-signaling) in the circulation and multiple organs, but most prominently the liver. Inhibition of IL-6 trans-signaling, using sgp130Fc transgenic mice or via specific genetic deletion of IL-6Ra on neutrophils, conferred protection against both diseases. Taken together, our findings suggest that a maternal HFD induces neonatal LGSI that predisposes to sLRI and subsequent asthma via neutrophil-mediated IL-6 trans-signaling.

## Introduction

Respiratory syncytial virus (RSV)-associated bronchiolitis is a significant source of morbidity, and a leading cause of death among infants and young children (Nair, Nokes et al. 2010). Additionally, prospective birth cohort and population studies have shown that frequent or severe lower respiratory tract infections (sLRIs) in early life increase the risk of developing childhood asthma (James, Gebretsadik et al. 2013). Whilst the causal mechanisms remain poorly understood, the prevailing view is that the aberrant immune response and associated tissue damage that occur as a consequence of the sLRI affects the early postnatal phase of immune and pulmonary development, laying the foundations that predispose to the onset of asthma (Jartti and Gern 2017, Lynch, Sikder et al. 2017). However, it is increasingly apparent that the status of the immune response at homeostasis heavily influences the host response, the course of infection, and downstream sequelae (Habibi, Thwaites et al. 2020).

Risk factors for sLRI include antibiotic use, never breastfeeding, high maternal body mass index, and maternal obesity (Håberg, Stigum et al. 2009, Murk, Risnes et al. 2011, Forno, Young et al. 2014). The latter has been associated with increased illness and disease risk in the infant, including recurrent wheeze, prolonged cough, infection severity, and asthma (Guerra, Sartini et al. 2013, Leermakers, Sonnenschein-van der Voort et al. 2013, Forno, Young et al. 2014, Rajappan, Pearce et al. 2017). As the incidence of adult obesity is increasing globally as a consequence of Western-style diets that are high in saturated fats, together with decreasing levels of physical activity (Ng, Fleming et al. 2014, Hruby and Hu 2015, Guthold, Stevens et al. 2018), this will inevitably have a negative impact on infant health and development. Notably, a poor maternal diet increases the risk of sLRI, as illustrated by a large population study which found that a maternal diet high in carbohydrates and low in fruit and vegetables was dose-dependently associated with severe bronchiolitis in the infant (Ferolla, Hijano et al. 2013). The consumption of diets high in fat leads to microbial dysbiosis (Murphy, Velazquez et al. 2015) and alters the metabolic activity of the host (Astrup, Dyerberg et al. 2008), leading to a state of low-grade systemic inflammation (LGSI) (Furman, Campisi et al. 2019). LGSI predisposes to an array of metabolic and chronic inflammatory diseases such as metabolic dysfunction-associated steatotic liver disease (Bhattacharjee, Kirby et al. 2017), cardiovascular disease (Ferrucci and Fabbri 2018), and lung diseases (Bennett, Reeves et al. 2018), and is typically characterized by increased baseline levels of cytokines, chemokines, and acute-phase proteins in serum, together with increased numbers of circulating myeloid cells (Mathur and Pedersen 2008, León-Pedroza, González-Tapia et al. 2015). However, the cellular and molecular processes that give rise to LGSI in infants remain poorly defined (Furman, Campisi et al. 2019).

The expression of IL-17A, a type-17 effector cytokine, is commonly elevated in the serum as well as multiple organs and tissues as a consequence of LGSI (Miossec and Kolls 2012). IL-17A is produced by various lymphocytes, including CD4^+^ T helper 17 cells, γδ-T cells and group 3 innate lymphocyte cells (ILC3s) (Park, Li et al. 2005, Ward and Umetsu 2014, Ullah, Revez et al. 2015), and the differentiation of these T cell populations is promoted by the combined actions of IL-6, IL-1β, and IL-23 (Manel, Unutmaz et al. 2008). Additionally, however, we and others have shown that IL-6 trans-signaling, mediated via an IL-6/soluble IL-6 receptor (sIL-6R) complex that directly activates gp130, even on cells without IL-6R expression, increases T cell IL-17A production (Ullah, Revez et al. 2015, Nadeem, Ahmad et al. 2020). Similar to IL-17A, systemic IL-6 and sIL-6R expression is elevated in obese people (van Bussel, Schouten et al. 2011, van Bussel, Henry et al. 2012, Kraakman, Kammoun et al. 2015) and this phenotype is recapitulated in adult mice fed a HFD (Xu, Pereira et al. 2017), consistent with the notion that IL-6 trans-signaling is pro-inflammatory (Rose-John, Jenkins et al. 2023). However, whether a maternal HFD promotes LGSI in the offspring remains to be determined, as does the contribution of IL-6 trans-signaling to this phenomenon and the role of neonatal LGSI in predisposing to sLRI and subsequent asthma. Here, using a high-fidelity preclinical model, we demonstrate that a maternal HFD promotes LGSI in the infant and predisposes to sLRI and subsequent asthma. LGSI was associated with increased systemic and hepatic levels of sIL-6R, IL-6, and IL-17A, and increased neutrophil numbers in the neonatal liver. Inhibition of IL-6 trans-signaling or the specific deletion of IL-6R on neutrophils ablated basal sIL-6R and IL-17A levels and conferred protection against sLRI and subsequent asthma. Taken together, our findings demonstrate that consequent to microbial dysbiosis in the neonatal gut, a maternal HFD promotes inflammatory granulopoiesis in the neonatal liver, leading to enhanced IL-6 trans-signaling, which in turn, predisposes to sLRI and childhood asthma.

## Results

### A maternal high fat diet predisposes the offspring to a sLRI

Breeding-age female and male mice were fed a high fat diet (HFD) or control fat diet (CFD) for three weeks prior to a timed mating, and throughout gestation and lactation (Fig. 1 A). The neonatal pups were inoculated with PVM (pneumonia virus of mice), the murine homologue of human RSV, at postnatal day (PND)7 (unless otherwise stated). Ingestion of the HFD increased the body weight of the dams prior to conception and during gestation but did not affect the weight of the offspring at PND7 (Fig. S1 A; and Fig. 1 B, left panel). However, in response to inoculation with PVM, neonates reared to HFD-fed mothers (hereafter referred to ‘HFD-reared pups’) exhibited stunted weight gain from 6 days post infection (dpi), as compared to CFD-reared neonates (Fig. 1 B, right panel). This was associated with increased viral load and delayed clearance, as assessed by quantifying PVM-positive airway epithelial cells (AECs) – a faithful measure of viral load (Lynch, Werder et al. 2016), impaired lung IFN-λ production, increased numbers of mucus-secreting AECs, and increased airway smooth muscle (ASM) area, indicative of airway remodeling (Fig. 1, C-F). Although epithelial sloughing can be a pathological feature of sLRI (Sikder, Rashid et al. 2023), it was not observed in the infected HFD-reared neonates (data not shown). Stunted weight gain is indicative of increased inflammation and disease severity, and consistent with this, there was a near 3-fold increase in lung neutrophil numbers at 5 dpi, and a doubling of eosinophil numbers at 7 dpi in HFD-compared to CFD-reared pups (Fig. 1 G; and Fig. S1, B and C). In addition to granulocytes, natural killer (NK) cells, group 1 innate lymphoid cells (ILC1s), ILC2s, ILC3s, as well as CD4^+^ T helper 1 (Th1), Th2, and Th17 cells, were all elevated in HFD-compared to CFD-reared pups (Fig. S2, A-C). Together, these findings indicated that a maternal HFD predisposes the offspring to a sLRI characterised by delayed viral clearance, structural changes to the lung architecture, and a hyper-inflammatory cellular response.

**Figure 1.**
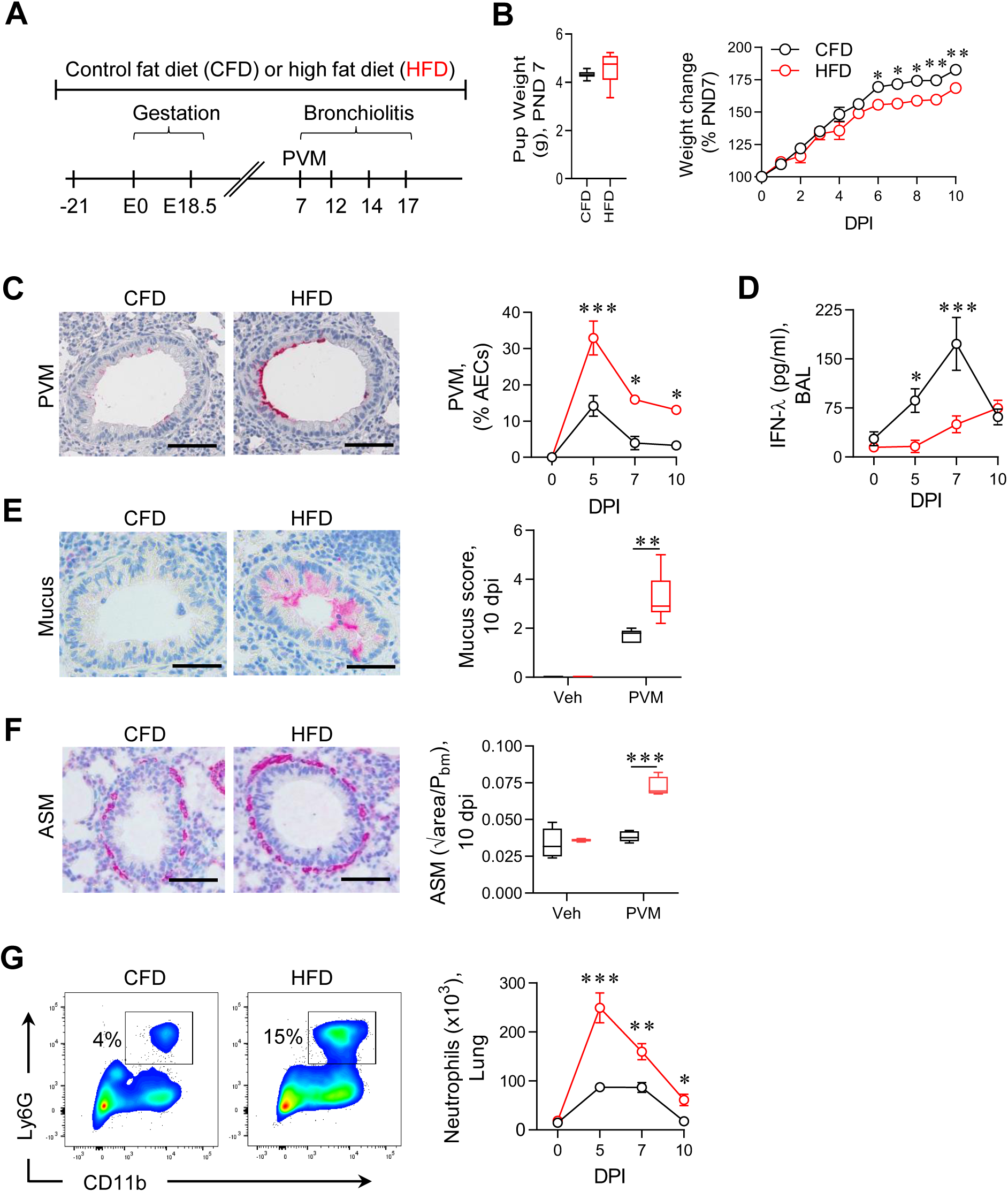
A maternal high fat diet predisposes the offspring to a sLRI. **(A)** Study design. **(B)** Bodyweight of pups at postnatal day (PND)7 (left panel) and weight change post PVM inoculation, expressed as percentage of weight at PND7. **(C)** Representative images of PVM immunoreactive (pink coloration) airway epithelial cells (AECs), ×20 magnification, scale bar = 100 µm (left panel) and quantification of PVM-positive AECs (right panel). **(D)** Interferon (IFN)-λ in the bronchoalveolar lavage (BAL) fluid. **(E)** Representative histology images of muc5ac (pink) in the airway, ×20 magnification, scale bar = 100 µm (left panel) and Mucus score. **(F)** Representative histology images of airway smooth muscle (ASM) surrounding the bronchioles (pink), ×20 magnification, scale bar = 100 µm (left panel) and ASM area. **(G)** Representative flow plots of neutrophils at 5 dpi (left panel) and neutrophils in the lung. Data are shown as mean ± SEM or box and whisker plots (median, quartiles, and range). Data are pooled from 2 independent experiments, *n* = 3-4 mice per group in each experiment. *P* values (*p< 0.05, **p< 0.01, ***p< 0.001) were derived by two-way ANOVA with Tukey’s post hoc test or Mann Whitney *U* test.

### HFD-reared pups develop mixed type-2/type-17 inflammation upon infection and low-grade systemic inflammation (LGSI) at steady-state

Given the elevated numbers of ILCs and T cells associated with type-1, type-2 and type-17 immunity, we next assessed the cytokine milieu in the airways. Despite the elevated numbers of NK cells, ILC1s, and Th1 cells, the level of IFN-γ was attenuated in the HFD-reared pups. By contrast, the type-2 associated cytokines, IL-4 and IL-5, together with the archetypal type-17 cytokine, IL-17A, were elevated in the airways of HFD-compared to CFD-reared pups (Fig. 2 A; and Fig. S3 A, full time course). This response was associated with increased airway expression of the instructive cytokines that promote type-2 immunity (i.e. IL-33, IL-25 and TSLP), as well as those that promote type-17 immunity (i.e. IL-23p19, IL-1β, IL-6) (Veldhoen, Hocking et al. 2006), including sIL-6R, which we have previously shown to increase IL-17A expression via IL-6 trans-signaling (Ullah, Revez et al. 2015) (Fig. 2 B; and Fig. S3 B, full time course). Unexpectedly, given the attenuated IFN-γ levels, expression of the Th1 differentiation factor IL-12p70, was also greater in the HFD-reared pups (Fig. 2 B). As a maternal HFD can promote LGSI in the offspring (Furman, Campisi et al. 2019), we next questioned whether the inflammatory phenotype was evident prior to the infection. Measuring the cytokines at PND7 revealed that all the instructive cytokines were markedly elevated at baseline in serum whereas, of the effector cytokines, only IL-17A was increased (Fig. 2, C and D). To determine whether IL-17A predisposed to sLRI, we inoculated HFD-reared IL-17A-deficient pups with PVM, and found that these mice were protected against stunted weight gain and virus-induced pathology (Fig. S4, A-F) Furthermore, the magnitude of inflammation was similar to that observed in infected CFD-reared pups. Of note, the absence of IL-17A decreased the elevated type-2 cytokine response, and de-repressed IFN-γ expression (Fig. S4 F). Collectively, these findings demonstrated that a maternal HFD profoundly affects the neonatal immune development as shown by increasing type-2 and type-17 instructive and effector cytokine expression, suppressing the expression of the type-1 cytokine IFN-γ, and that as a consequence the offspring is predisposed to a sLRI in an IL-17A-dependent manner.

**Figure 2.**
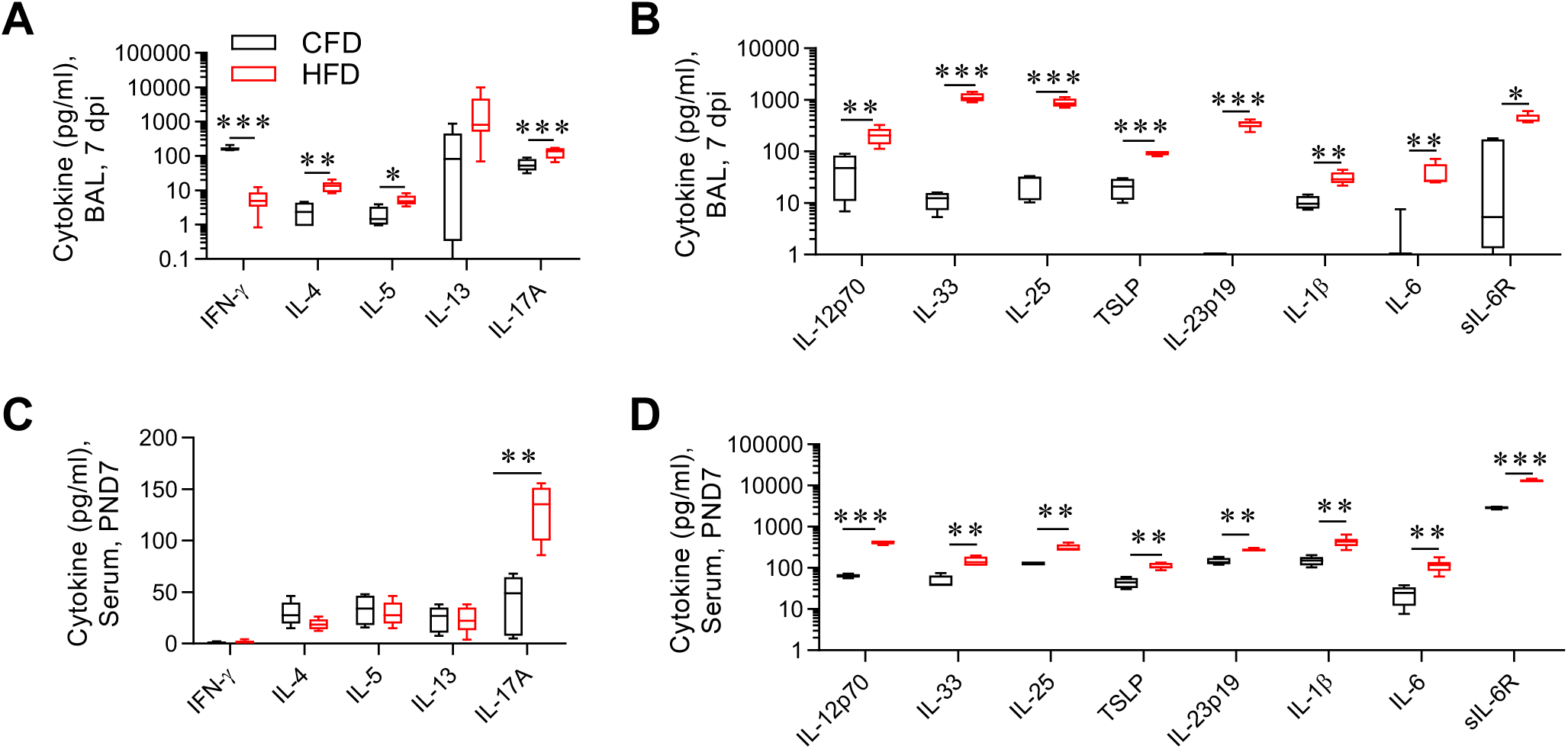
HFD-reared pups develop mixed type-2/type-17 inflammation upon infection and LGSI at steady-state. **(A)** IFN-γ, IL-4, IL-5, IL-13 and IL-17A and **(B)** IL-12p70, IL-33, IL-25, Thymic stromal lymphopoietin (TSLP), IL-23p19, IL-1β, IL-6 and sIL-6R in the BAL fluid at 7 dpi. **(C)** IFN-γ, IL-4, IL-5, IL-13, IL-17A and **(D)** IL-12p70, IL-33, IL-25, TSLP, IL-23p19, IL-1β, IL-6 and sIL-6R in serum at PND7. Data are shown as box-and-whisker plots (median, quartiles, and range). Data are pooled from 2 independent experiments, *n* = 3-4 mice per group in each experiment. *P* values (*p< 0.05, **p< 0.01, ***p< 0.001) were derived by Mann Whitney *U* test.

### HFD-reared pups that develop sLRI are predisposed to subsequent asthma

Severe RSV bronchiolitis in infancy is a major risk factor for the development of childhood asthma (James, Gebretsadik et al. 2013). To determine whether the maternal HFD and increased LRI severity in the HFD-reared pups would predispose to experimental asthma, we exposed PVM-or diluent-inoculated mice to low-dose cockroach allergen extract (CRE) at PND42, 49, 56 and 63 (Fig. 3 A), an approach that we have described previously (Lynch, Werder et al. 2016, Sikder, Rashid et al. 2023). The HFD-reared mice gained more weight than their CFD counterparts after weaning, and consistent with the human epidemiology, the HFD-reared mice that developed a sLRI in infancy developed the hallmark features of asthma in later life (upon exposure to CRE), including increased ASM area, mucus hyper-production and increased granulocytic inflammation (Fig. 3, B and C; and Fig. S5, A and B). In contrast, PVM/CRE-exposed CFD-reared mice did not develop experimental asthma, nor CFD- or HFD-reared pups exposed to either PVM or CRE alone. Similar to the inflammatory phenotype observed during the sLRI, PVM/CRE-exposed HFD-reared pups expressed elevated levels of type-2 and type-17 effector cytokines, whereas the production of IFN-γ was suppressed (Fig. 3 D; and Fig. S5 C). However, the type-17 response appeared more dominant in the asthma phase, as demonstrated by the prevalence of CD4^+^ Th17 cells (relative to CD4^+^ Th2 cells, ILC2, or ILC3), high IL-17A levels, and a bias of the granulocyte compartment towards neutrophils (Fig. 3, C-E; and Fig. S5 D). Thus, the modules of inflammation that developed in early-life in response to the acute infection were largely recapitulated in response to allergen exposure in later-life, although the latter appeared to be dominated by Th17 cells rather than ILC3s.

**Figure 3.**
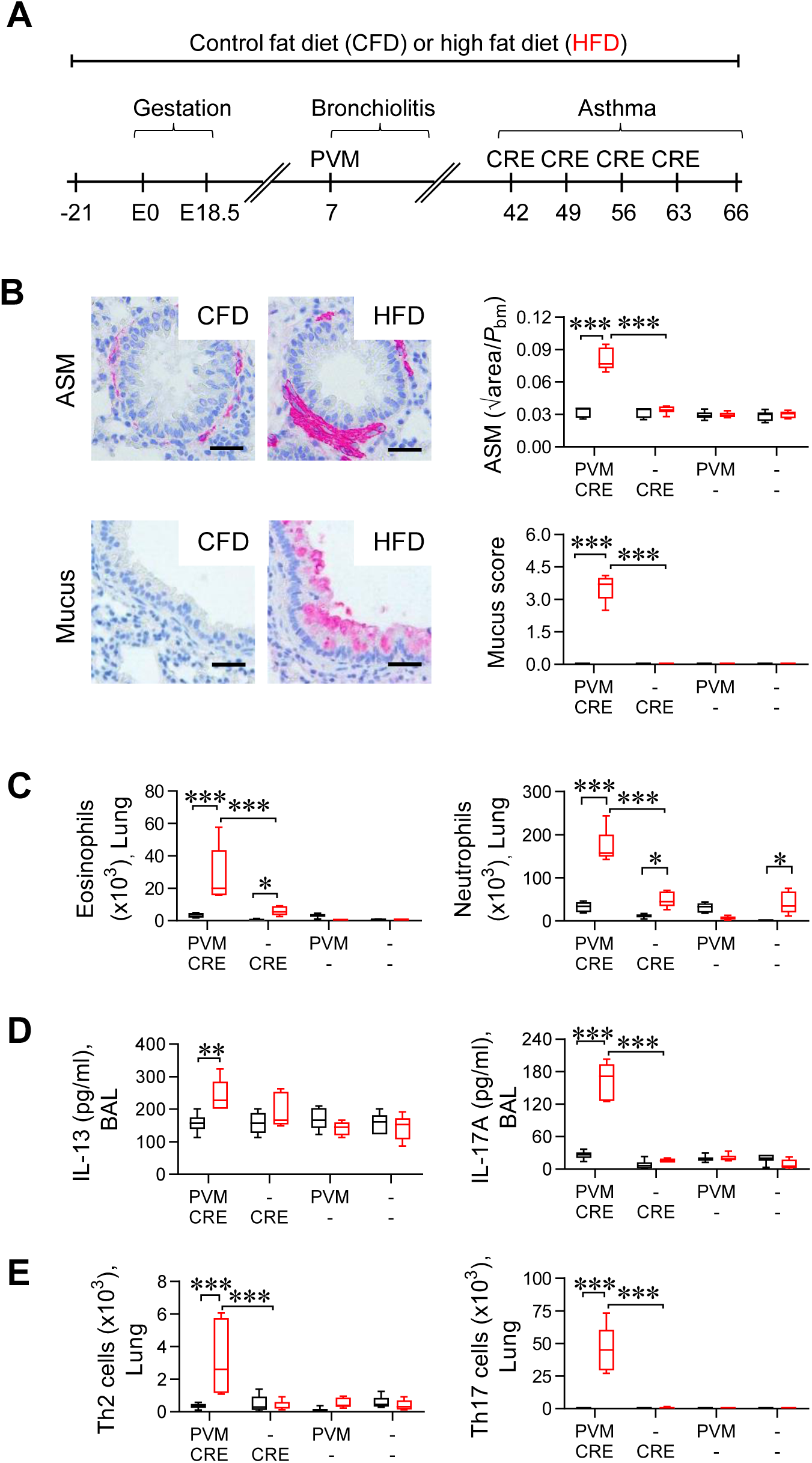
HFD-reared pups that develop sLRI in early life are predisposed to subsequent asthma. **(A)** Study design. **(B)** Representative histology images of ASM surrounding the bronchioles (pink), ×20 magnification, scale bar = 100 µm (top left panel) and ASM score (top right panel); Representative histology images of muc5ac (pink) in the airway, ×20 magnification, scale bar = 100 µm (bottom left panel) and Mucus score (bottom right panel). **(C)** Eosinophils (left panel) and neutrophils in the lung. **(D)** IL-13 and IL-17A expression in BAL fluid. **(E)** CD4^+^ T helper type-2 (Th2) (left panel) and Th17 cells in the lung. Data are shown as box-and-whisker plots (median, quartiles, and range). Data are pooled from 2 independent experiments, *n* = 3-4 mice per group in each experiment. *P* values (*p< 0.05, **p< 0.01, ***p< 0.001) were derived by two-way ANOVA with Tukey’s post hoc test.

### Blockade of IL-6 trans-signaling protects HFD-reared mice against sLRI and asthma

As HFD-reared IL-17A-deficient pups were protected from developing sLRI, we sought to address the molecular events by which a maternal HFD promotes high systemic levels of IL-17A. As we and others had previously shown that IL-6 trans-signaling, through the combined actions of IL-6 and sIL-6R promotes IL-17A production by T cells (Jones, McLoughlin et al. 2010, Ullah, Revez et al. 2015), we hypothesised that the blockade of IL-6 trans-signaling using sgp130Fc transgenic mice, where sgp130Fc-protein is genetically overexpressed to prevent gp130 activation by the IL-6/sIL-6R complex, would protect against sLRI and the subsequent development of asthma. Of note, the sgp130Fc protein does not affect IL-6 signaling via the membrane bound IL-6R (Jostock, Müllberg et al. 2001). Prior to infection, the serum concentration of sIL-6R, IL-6, and IL-23p19, did not differ between HFD-reared WT and HFD-reared sgp130Fc transgenic pups (Fig. 6 A). However, serum levels of IL-1β and IL-17A were significantly lower in the HFD-reared sgp130Fc transgenic pups (Fig. 4 A; and Fig. S6 A). In response to PVM inoculation, the levels of IL-23p19, IL-1β, but not sIL-6R or IL-6 were significantly lower in the airways of HFD-reared sgp130Fc transgenic pups compared to HFD-reared WT pups (Fig. S6 B). This was associated with a delayed increase in IL-17A expression, an attenuated IL-4 response, the restoration of IFN-γ expression in the airways, and decreased numbers of ILC2s, ILC3s and neutrophils in the lungs (Fig. 4, B-D). Consistent with the observed immunological changes, inhibition of IL-6 trans-signaling in HFD-reared pups prevented body weight stunting and attenuated the development of mucus hyperproduction and ASM remodeling (Fig. 4, E-G; and Fig. S6 C). Unexpectedly, viral load was not decreased in HFD-reared sgp130Fc transgenic pups compared to HFD-reared pups at 5 dpi, however, there were fewer PVM-immunoreactive AECs at 7 dpi, and this was associated with the restoration of IFN-λ production (Fig. 4, H and I). These findings suggested that blockade of IL-6 trans-signaling would break the nexus between sLRI and later-life susceptibility to asthma. As postulated, HFD-reared sgp130Fc transgenic mice were protected from developing experimental asthma, as shown by the absence of mucus hyperproduction, ASM remodeling, and granulocytic inflammation compared to their wildtype control counterparts (Fig. 4, J and K). Notably, the attenuated immunopathology was associated with a switch from type-2/17 to type-1 immunity (Fig. 4 L). Collectively, these findings highlight that a maternal HFD increases the infant’s susceptibility to LGSI, sLRI, and allergic asthma through enhanced activation of IL-6 trans-signaling.

**Figure 4.**
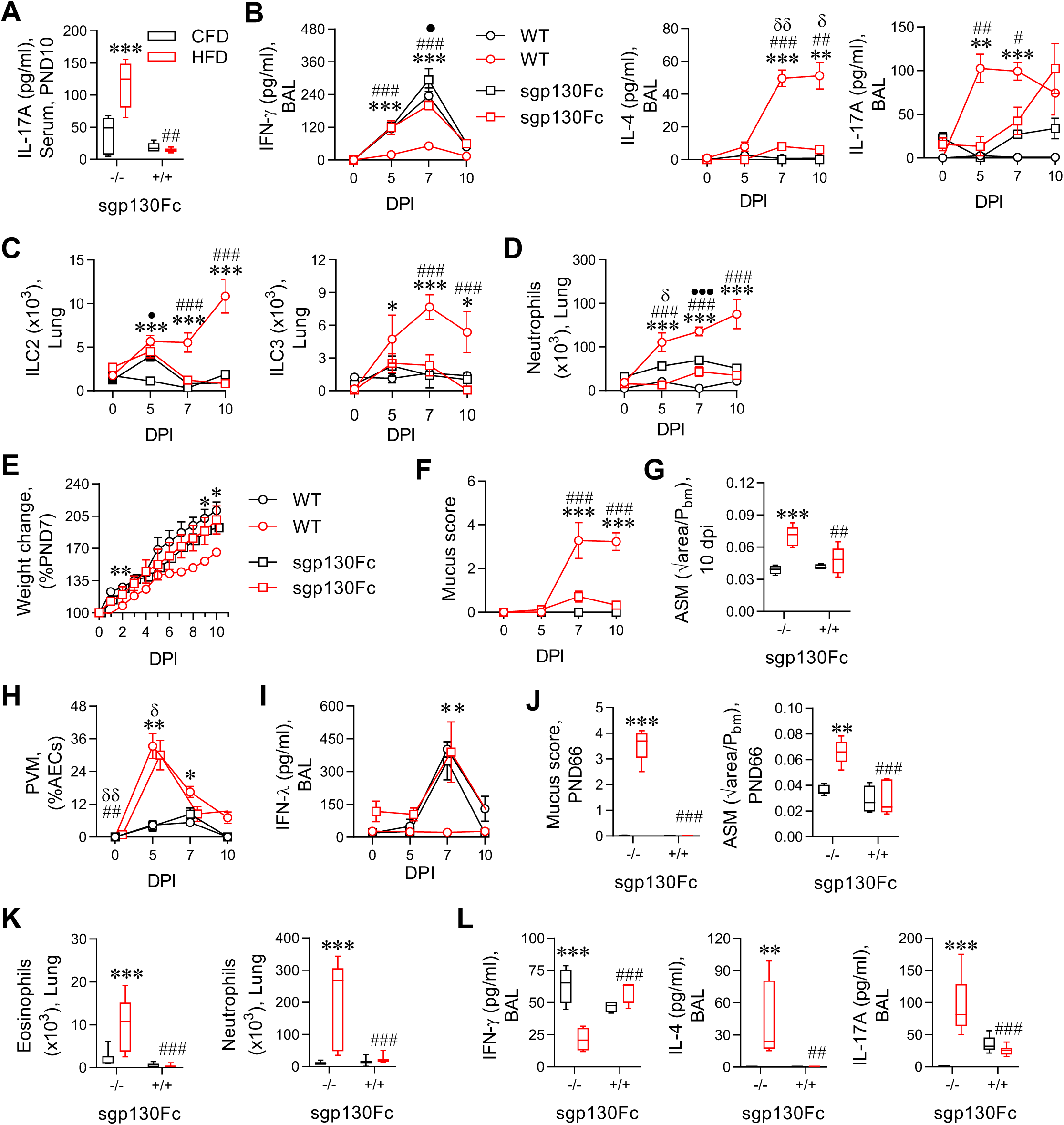
Blockade of IL-6 trans-signaling protects HFD-reared mice against sLRI and asthma. **(A)** IL-17A in the serum at PND10. **(B)** IFN-γ, IL-4 and IL-17A levels in BAL fluid post PVM infection. **(C)** Group-2 innate lymphoid cells (ILC2s) and ILC3s. **(D)** Neutrophils in the lung. **(E)** Weight change post PVM inoculation, expressed as percentage of weight at PND7. **(F)** Mucus score. **(G)** Airway smooth muscle (ASM) area at 10 dpi. **(H)** Viral load. **(I)** IFN-λ levels in BAL fluid. **(J-L)** Asthma phase: (J) Mucus score and ASM area. (K) Eosinophils and neutrophils in the lung. (L) IFN-γ, IL-4 and IL-17A levels in BAL fluid. Data are shown as mean ± SEM or box- and-whisker plots (median, quartiles, and range). Data are pooled from 2 independent experiments, *n* = 3-4 mice per group in each experiment. *P* values (*p< 0.05, **p< 0.01, ***p< 0.001) were derived by two-way ANOVA with Tukey’s post hoc test. * were used to compare CFD_WT vs HFD_WT, # used to compare HFD WT vs HFD sgp130Fc tg mice, δ used to compare CFD_sgp vs HFD_ sgp130Fc tg mice, • used to compare CFD_ sgp130Fc tg mice vs CFD_WT.

### The maternal HFD influences infant gut microbiome composition and predisposes to sLRI

To elucidate the pathogenic mechanisms by which a maternal HFD predisposes to enhanced IL-6 trans-signaling and sLRI in early-life, we next assessed whether the cross-fostering of pups, within 24 hours of birth, would affect susceptibility. Switching CFD-reared (*in utero*) pups to HFD-fed dams (C->H) predisposed them to sLRI, as demonstrated by stunted weight gain, increased viral load, diminished IFN-λ production, increased ASM area, and a switch from type-1 to type-2/ type-17 inflammation (Fig. 5, A-G; and Fig. S7, A-D). In contrast, the nursing of HFD-reared (*in utero*) pups by CFD-fed dams conferred protection against sLRI (Fig. 5, A-G; and Fig. S7, A-D), indicating that the predominant impact of the maternal diet is elicited postnatally and suggesting a potential role of the maternal microbiome, which can be vertically transmitted and modified by the milk (Sikder, Rashid et al. 2023). To determine whether the maternal HFD affects the composition of the mother and infant gut microbiome, we performed 16S rRNA gene amplicon sequencing on fecal pellets. As expected, the maternal HFD significantly altered the composition of bacterial communities associated with the feces of dams at partum (*p =* 0.012, PERMANOVA) and infants at PND7 (*p =* 0.03, PERMANOVA) (Fig. 5h). Indicator analysis revealed that a *Streptococcus* (Otu 6) and a *Muribacter* (OTU 192) population were significantly positively associated with the CFD (*P* < 0.025), while a representative of the *Lactobacillus* was positively associated with the HFD (*P* = 0.029). Despite the altered microbial composition, epithelial barrier permeability was unaffected in the neonatal mice (Fig. 5 I), suggesting that the observed immunological phenotypes were not a consequence of ‘leaky gut’. To address whether the altered neonatal microbiome contributes to disease susceptibility, we next performed fecal microbiome transplants (FMT) using fecal material obtained from CFD- or HFD-reared pups (Fig. S7 E). Transfer via oral gavage of a fecal slurry from HFD-reared pups to CFD-reared pups (i.e., H(F)->C) was sufficient to predispose to sLRI, as shown by stunted weight gain, increased immunopathology, and a switch in the inflammatory responses to levels similar to those observed in HFD-reared pups (Fig. 5, J-N). In contrast, an FMT using material from CFD-reared pups conferred protection to HFD-reared pups (C(F)->H). Taken together, these findings demonstrated that the maternal HFD affects the composition of the neonatal gut microbiome, which in turn predisposes to sLRI.

**Figure 5.**
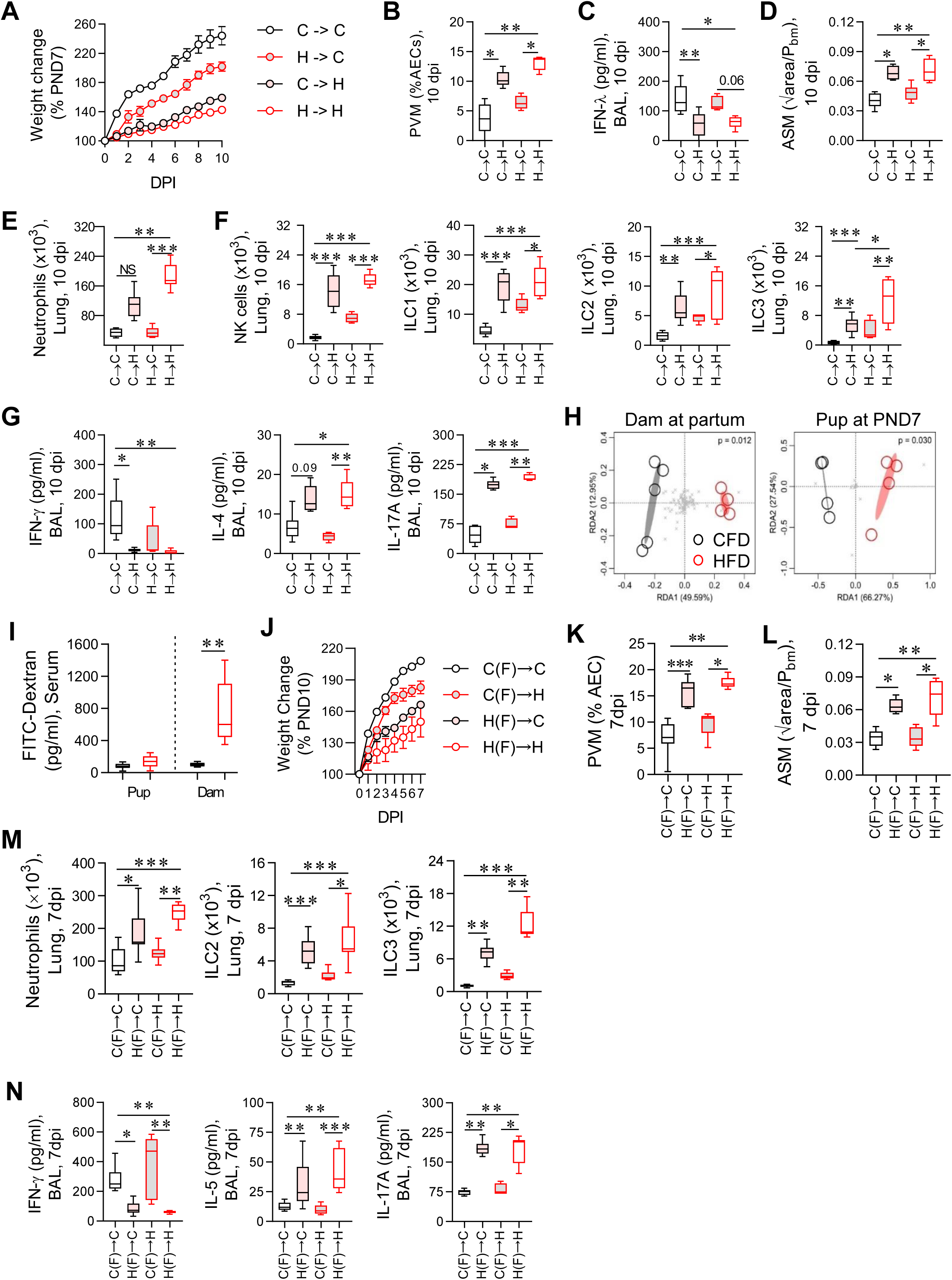
The maternal HFD influences infant gut microbiome composition and predisposes to sLRI. Pups were cross fostering mothers consuming a CFD to a HFD (C→H), from a HFD to a HFD (H→H), from a CFD to a CFD (C→C) and from a HFD to a CFD (H→C). **(A)** Weight change post PVM inoculation, expressed as percentage of weight at postnatal day (PND)7. **(B)** Viral load. **(C)** IFN-λ in BAL fluid at 10 dpi. **(D)** Airway smooth muscle (ASM) area at 10 dpi. **(E)** Lung neutrophils. **(F)** Natural killer (NK) cells, ILC1s, ILC2s and ILC3s in the lung at 10 dpi. **(G)** IFN-γ, IL-4, IL-17A in BAL fluid at 10 dpi. **(H)** An RDA ordination highlighting the effect of diet on the differences in bacterial community composition between fecal samples collected from mothers consuming a HFD or a CFD at E0 (P<0.012, PERMANOVA) and neonates at PND7 (P<0.03, PERMANOVA). Each circle represents a sample, and the ellipses represent the standard deviation around the centroid of each group. (**I)** FITC-Dextran level in the serum of the females and their neonates. **(J-N)** Fecal microbiota transplant (FMT) experiment (Study design shown in Fig. S7 E). FMT from CFD-reared pups to HFD-reared neonates [C(F)→H]; from HFD-reared pups to CFD-reared neonates [H(F)→C]; from CFD-reared pups to CFD-reared neonates [C(F)→C]; from HFD-reared pups to HFD-reared neonates [H(F)→H]. (J) Weight change post PVM inoculation, expressed as percentage of weight at PND7. (K) Viral load. (L) Airway smooth muscle (ASM) area. (M) Neutrophils, ILC2s and ILC3s in lungs. (N) IFN-γ, IL-5 and IL-17A levels in BAL fluid at 7 dpi. Data are shown as mean ± SEM or box-and-whisker plots (median, quartiles, and range). Data are pooled from 2 independent experiments, *n* = 3-4 mice per group in each experiment. *P* values (*p< 0.05, **p< 0.01, ***p< 0.001) were derived by one-way or two-way ANOVA with Tukey’s post hoc test.

**Figure 6.**
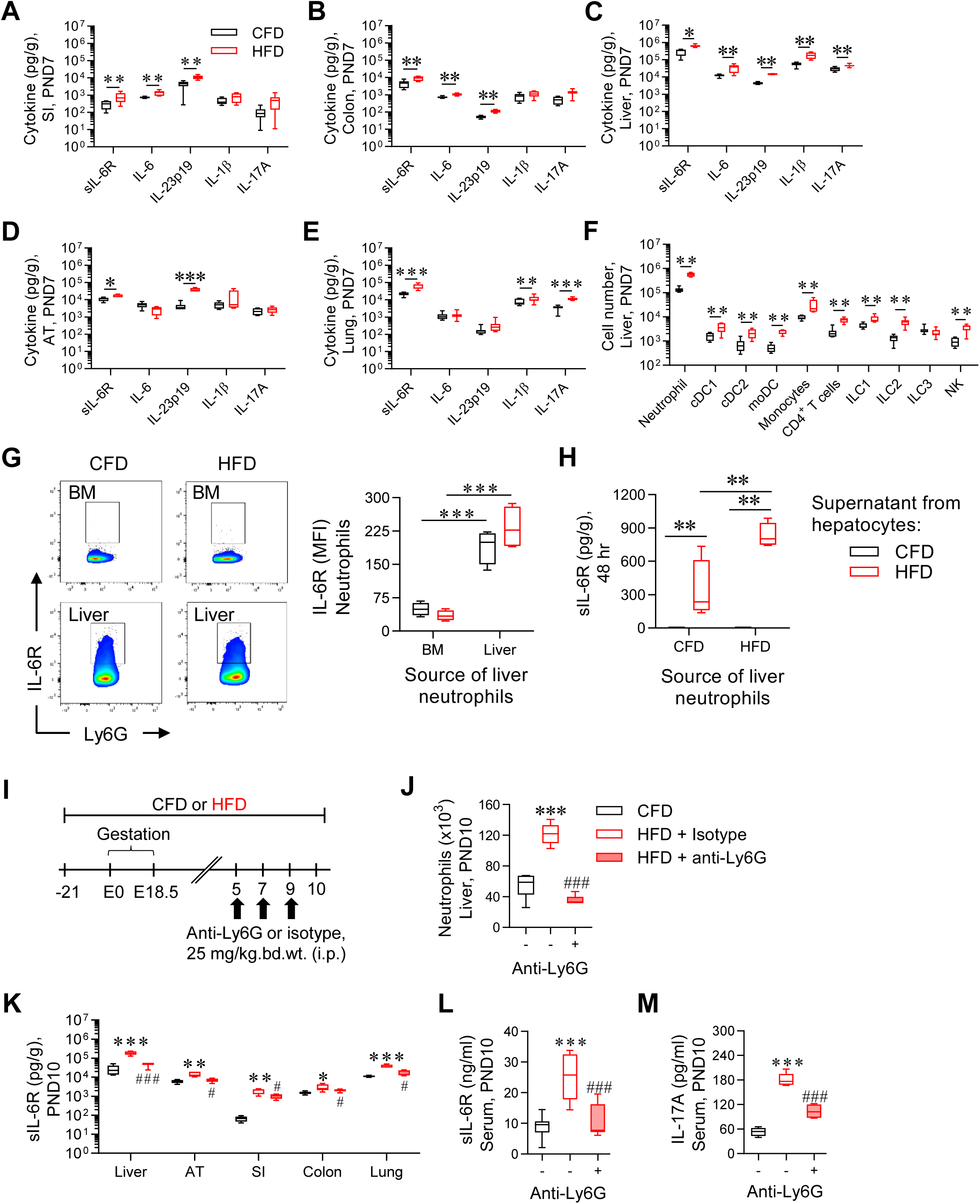
Neutrophil depletion ameliorates the elevated IL-6 trans-signaling in HFD-reared pups. **(A-E)** sIL-6R, IL-6, IL-23p19, IL-1β and IL-17A levels in the (A) small intestine (SI), (B) colon, (C) liver, (D) adipose tissue (AT) and (E) lungs. **(F)** IL-6R-expressing hematopoietic cells in the liver. **(G)** Representative flow plot of IL-6R expression on neutrophils in the bone marrow (BM) and liver and mean fluorescent intensity (MFI) of IL-6R. **(H)** FACS-sorted liver neutrophils cultured with the supernatant of primary hepatocytes purified from CFD- and HFD-reared neonates. **(I)** Study design of neutrophil depletion. **(J)** Neutrophils in the liver. **(K)** sIL-6R level in the liver, AT, SI, colon, and lungs. **(L)** sIL-6R and **(M)** IL-17A levels in serum. Data are shown as box-and-whisker plots (median, quartiles, and range). Data are pooled from 2 independent experiments, *n* = 3-4 mice per group in each experiment. *P* values (*p< 0.05, **p< 0.01, ***p< 0.001) were derived by one-way ANOVA with Tukey’s post hoc test or Mann Whitney *U* test. * were used to compare CFD vs HFD + Isotype and # used to compare HFD + Isotype vs HFD + anti-Ly6G.

### Neutrophil depletion ameliorates the elevated IL-6 trans-signaling in HFD-reared pups

Having established that increased susceptibility to sLRI is associated with an altered gut microbial composition and a state of LGSI - most notably enhanced type-17 inflammation - we next sought to identify the source of IL-6 trans-signaling by assessing key organs and tissues known to be affected by a HFD. The expression of sIL-6R and IL-23p19 was elevated in HFD-compared to CFD-reared pups in all the sites analysed, namely the small intestine, colon, inguinal adipose tissue, and liver (Fig. 6, A-E). IL-1β was elevated in the liver and lung only, and IL-6 was increased in the small intestine, colon, and liver, although the concentration in the gut was a log lower than that of the liver. Taken together, these findings identified the liver as a rich source of both sIL-6R and IL-6 production, and implicated the liver as a probable site of active IL-6 trans-signaling, Consistent with this, IL-17A levels were elevated in the liver (and lung), but not the small intestine, colon, or adipose tissue of HFD-reared pups (Fig. 6, A-E). Further support of a key role for the liver stemmed from changes in sIL-6R, IL-6, and IL-23p19 levels in the liver and serum in response to the FMT (Fig. S8, A-B) and these shifts in expression associated with LRI severity (Fig. 5). We hypothesised that the increase in liver sIL-6R expression derived from increased inflammatory cell expansion/recruitment to the liver. Remarkably, a maternal HFD increased the numbers of all immune cell types analysed in the neonatal liver, including neutrophils, dendritic cells, monocytes, T cells, ILCs, and NK cells, although neutrophils were by far the most prevalent (Fig. 6 F). Of note, flow cytometric analysis of neutrophil IL-6R revealed that the receptor was highly expressed on neutrophils in the liver, but not the bone marrow (Fig. 6 G), an effect that was independent of maternal consumption of dietary fat. To determine whether neutrophils produce sIL-6R, neutrophils were FACS-purified from the livers of CFD- and HFD-reared pups, and cultured with the supernatant (sIL-6R was undetectable; data not shown) collected from primary neonatal hepatocytes isolated from CFD- and HFD-reared pups. Remarkably, neutrophils stimulated with hepatocyte supernatant from CFD-reared pups failed to release sIL-6R. In contrast, stimulation with the hepatocyte supernatant of HFD-reared pups led to a marked increase in sIL-6R levels, and this response was even greater when the neutrophils were derived from the liver of HFD-reared pups (Fig. 6 H). Neutrophil numbers were also elevated in the SI, colon, and lungs, though not the bone marrow or adipose tissue (Fig. S8 C). To assess the relative contribution of neutrophils to the expression of sIL-6R, we treated the HFD-reared pups with anti-Ly6G or an isotype-matched control antibody (Fig. 6 I). Anti-Ly6G depleted liver neutrophil numbers and led to a marked decrease in sIL-6R expression in the liver (and other organs/tissues) and the serum, decreased liver IL-6 and IL-1β expression, and ablated the elevated serum IL-17A levels in HFD-reared pups (Fig. 6, J-M; and Fig. S8 D). Taken together, these data demonstrate that a maternal HFD promotes IL-6 trans-signaling in multiple neonatal organs, and particularly in the liver, and suggest that neutrophils are the primary source of sIL-6R in this context, predisposing to the development of LGSI.

### Genetic ablation of IL-6R on neutrophils ameliorates LGSI, LRI severity, and subsequent asthma

To definitively implicate neutrophil-derived IL-6R in the development of LGSI and predisposition to sLRI and later asthma, we crossed Ly6G^cre^ mice with IL-6Ra flox (IL-6Ra^fl/fl^) mice to specifically delete IL-6R expression by neutrophils. Flow cytometric analysis confirmed that the gene-deletion of IL-6R on liver neutrophils ablated neutrophil IL-6R expression in Ly6G^Cre^IL-6Ra^fl/fl^ but not control IL-6R^fl/fl^ pups (Fig. 7 A). Genetic ablation of IL-6R on neutrophils halved the number of liver neutrophils in HFD-reared Ly6G^Cre^IL-6Ra^fl/fl^ pups compared to the IL-6Ra^fl/fl^ controls at PND10, indicating that neutrophil-derived IL-6R contributes to the amplification of neutrophilic inflammation (Fig. 7 B). sIL-6R levels were markedly lower in the liver and other organs/tissues in LyG6^cre^IL-6Ra^fl/fl^ relative to control IL-6Ra^fl/fl^ mice (Fig. 7 C), further implicating neutrophils as the major source of IL-6R or suggesting that neutrophil-derived IL-6R plays a seminal role in the initiation and amplification of sIL-6R production by other cell types. In accordance with IL-6 trans-signaling promoting LGSI, the levels of IL-6, IL-1β, IL-23 and IL-17A were ablated in multiple organs, tissues and the circulation of HFD-reared Ly6G^Cre^IL-6Ra^fl/fl^ pups (Fig. 7 D; and Fig. S9 A). Accordingly, we hypothesised that HFD-reared Ly6G^Cre^IL-6Ra^fl/fl^ pups would be protected against sLRI, and indeed, genetic ablation of IL-6R on neutrophils prevented the stunted weight gain observed in HFD-reared IL-6Ra^fl/fl^ pups and HFD-reared WT pups, decreased the viral load, increased IFN-λ expression, and prevented the development of airways remodeling (Fig. 7, E-H and Fig. S9 B). This was associated with a marked decrease in lung neutrophils, eosinophils, all ILC subsets, type-2/type-17 cytokines, and the restoration of IFN-γ production (Fig. 7, I and J; and Fig. S9, C-F). Lastly, we assessed whether the gene-deletion of neutrophil IL-6R expression would reverse the susceptibility of HFD-reared pups to develop later asthma. As expected, in light of the mild LRI that developed in PVM inoculated HFD-reared Ly6G^Cre^IL-6Ra^fl/fl^ pups, these mice were protected against the later development of CRE-induced asthma (Fig. 7, K and L). This effect was once again associated with a shift from a mixed type-2/17 to a type-1 immune response (Fig. 7 M; and Fig. S9, G and H).

**Figure 7.**
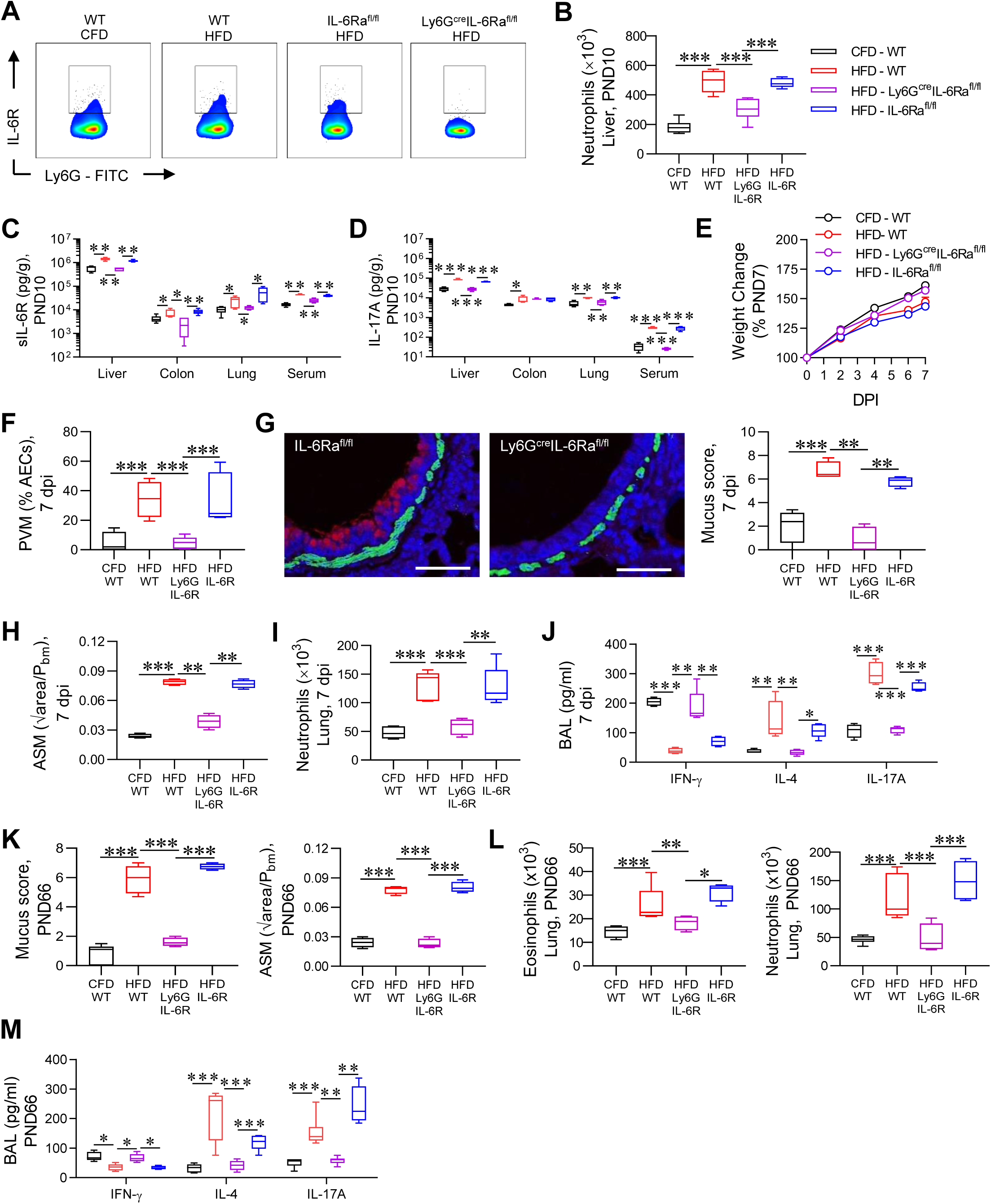
Genetic ablation of IL-6R on neutrophils ameliorates LGSI, LRI severity, and subsequent asthma. **(A)** Representative flow plot of IL-6R expression on neutrophils from WT, Ly6G^cre^IL-6Ra^fl/fl^ and IL-6Ra^fl/fl^ mice. **(B)** Liver neutrophils at PND10. **(C)** sIL-6R and **(D)** IL-17A in the liver, colon, lungs and serum. **(E)** Weight change post PVM inoculation, expressed as percentage of weight at PND7. **(F)** Viral load. **(G)** Representative immunofluorescence image of muc5ac (red), ASM (green), ×20 magnification, scale bar = 50 µm and mucus score. **(H)** ASM area. **(I)** Neutrophils in the lung. **(J)** IFN-γ, IL-4 and IL-17A levels in BAL fluid. **(K-M)** Asthma phase: (K) Mucus score and ASM area at PND66. (L) Eosinophils and neutrophils in the lung. (M) IFN-γ, IL-4 and IL-17A in BAL fluid. Data are shown as mean ± SEM or box-and-whisker plots (median, quartiles, and range). Data are pooled from 2 independent experiments, *n* = 3-4 mice per group in each experiment. *P* values (*p< 0.05, **p< 0.01, ***p< 0.001) were derived by one-way or two-way ANOVA with Tukey’s post hoc test.

## Discussion

Here, we demonstrated that neonatal mice nurtured by a mother consuming a HFD develop a sLRI upon infection with PVM, and are predisposed to develop subsequent asthma upon challenge with cockroach allergen. Prior to the infection, HFD-reared pups presented with LGSI, characterized by the increased expression of multiple cytokines in various organs and the circulation, most notably IL-17A, and IL-6 and sIL-6R, which together implicated IL-6 trans-signaling-induced type-17 inflammation. Of note, blockade of IL-6 trans-signaling ameliorated some aspects of LGSI, namely IL-1β and IL-17A expression, and conferred protection against sLRI and subsequent asthma. Lastly, we identified mechanistically that maternal HFD-induced LGSI in the offspring, and subsequent susceptibility to sLRI and asthma, is mediated via neutrophil-derived sIL-6R.

In high-income nations, ∼60% of expectant mothers are either overweight or obese and similar numbers are predicted in lower-income nations by 2040. Significantly, maternal obesity or the consumption of a high-fat diet (HFD) increases the risk of severe RSV bronchiolitis in the infant (Håberg, Stigum et al. 2009, Ferolla, Hijano et al. 2013, Luhar, Timæus et al. 2020). To simulate the human epidemiology, we used a preclinical model and found that in response to PVM inoculation, HFD-reared pups developed a mixed type-2 and type-17 response and an attenuated IFN-γ response, despite the presence of high numbers of NK cells, ILC1s, and Th1 cells in the lungs. Blockade of IL-6 trans-signaling, either via sgp130Fc transgenic mice or the genetic deletion of neutrophil IL-6R expression, or genetic ablation of IL-17A, restored the expression of IFN-γ, demonstrating that the pro-IL-17A microenvironment suppressed type-1 immunity, and most likely contributed to the increased viral load in the lungs. Of note, sLRIs are commonly associated with low IFN-γ expression (Selvaggi, Pierangeli et al. 2014, Hillyer, Mane et al. 2017, Hijano, Vu et al. 2019), and obesity is known to suppress IFN-γ production (Michelet, Dyck et al. 2018). Consistent with the type-2/type-17 cytokine milieu, both eosinophils and neutrophils were elevated in the lungs of HFD-reared pups, although neutrophils predominated. Although airway remodeling was apparent, as indicated by the increased numbers of mucus-producing cells and ASM area, the integrity of the airway epithelium was not compromised, unlike in other preclinical models that we have characterized, where necroptosis is evident and contributes to immunopathology (Simpson, Loh et al. 2020, Sikder, Rashid et al. 2023). Similar to the asthma field, it is now increasingly recognized that there are different subtypes or phenotypes of bronchiolitis, with each likely to have a distinct endotype or pathogenic process (Polack, Stein et al. 2019). Understanding these underlying mechanisms and the identification of specific biomarkers is critical to developing new therapies to alleviate the immune-mediated pathology that underpins both acute infections and the subsequent development of chronic diseases, such as asthma or long COVID (Raita, Pérez-Losada et al. 2021, Carraro, Ferraro et al. 2022, Fujiogi, Zhu et al. 2022, Ooka, Raita et al. 2022). In the subtype of bronchiolitis simulated here, stemming from a maternal HFD, disease predisposition (underpinned by LGSI) and disease pathogenesis was mediated by IL-6 trans-signaling and the downstream production of IL-17A.

LGSI is a risk factor for multiple metabolic diseases (Furman, Campisi et al. 2019), and is commonly associated with increased expression of IL-17A (Miossec and Kolls 2012). In the HFD-reared pups, IL-17A levels were increased in the circulation and various organs at homeostasis, whereas the other archetypal T helper cytokines (e.g., IL-4, IL-5, IFN-γ) were unaffected prior to infection. In contrast, several innate cytokines, especially those known to promote ILC and T helper cell differentiation, were elevated both in the serum and in multiple organs and tissues. We focused on IL-23p19, IL-1β, IL-6, and sIL-6R, as these cytokines are implicated in promoting type-17 immunity, and IL-6 and sIL-6R in particular, since IL-6 trans-signaling is typically pro-inflammatory and promotes IL-17A production (Ullah, Revez et al. 2015, Nadeem, Ahmad et al. 2020), It is also noteworthy that sIL-6R levels are elevated in the serum of overweight children (McFarlin, Johnston et al. 2007), and adult mice consuming a HFD (Kraakman 2014, Timper, Denson et al. 2017, Xu, Pereira et al. 2017). We found that all four cytokines were highly expressed in the liver, and that this expression decreased upon a protective intervention (e.g. FMT), consistent with the notion that the observed phenotype was linked to microbial dysbiosis. Future studies should seek to further define the associations between the gut microbiome and the promotion of excessive liver granulopoiesis in HFD-reared pups. IL-6R is expressed on both structural cells, such as hepatocytes, and hematopoietic cells, in particular monocytes and neutrophils (McFarland-Mancini, Funk et al. 2010, Farahi, Paige et al. 2017, Yousif, Ronsard et al. 2021). However, in the liver of HFD-reared pups, neutrophil numbers were markedly increased, and greater than twenty times more prevalent than monocytes, implicating neutrophils as the predominant source of sIL-6R. Intriguingly, few if any bone marrow neutrophils expressed IL-6R whereas in the liver, irrespective of maternal diet, approximately 30% of neutrophils were positive, consistent with the notion that neutrophils respond and adapt to their local environment (Qu, Jin et al. 2023). Intriguingly, liver neutrophil-derived sIL-6R was generated following stimulation with the supernatant of cultured hepatocytes, but only when the hepatocytes were purified from HFD-reared neonates, suggesting that the hepatocytes were differentially activated. The factor(s) regulating neutrophil IL-6R expression would be a logical therapeutic target but at this stage, remain to be determined. Neutrophil depletion ablated the expression IL-1β, IL-23, and IL-6 in the liver, markedly lowered sIL-6R in all compartments analyzed and attenuated serum IL-17A levels. Critically, this phenotype was recapitulated when IL-6R was genetically deleted specifically from neutrophils, implicating a key role for neutrophils in promoting LGSI in neonatal mice as a consequence of a maternal HFD. Remarkably, in the absence of neutrophil IL-6R, a maternal HFD did not predispose the offspring to either a sLRI or experimental asthma, highlighting a key role of neutrophils in linking microbial dysbiosis to the development of LGSI and disease susceptibility, and suggesting that interventions that target IL-6 trans-signaling, such as olamkicept, which is showing clinical benefit in ulcerative colitis and Crohn’s disease (Schreiber, Aden et al. 2021, Zhang, Chen et al. 2023), may hold promise in specific endotypes of sLRI. Intriguingly, IL-6 trans-signaling has been implicated in type-2 low asthma, commonly associated with neutrophilic inflammation and elevated sputum IL-8 and IL-1β (Jevnikar, Ostling et al. 2019), although the underlying cause of the elevated sIL-6R remains unknown.

A limitation of our studies is that the cross-fostering and FMT were performed in the neonatal period prior to the infection, and the genetic interventions were constitutive rather than inducible. Hence, it is not possible to assess whether ablating the state of LGSI alone is sufficient to prevent sLRI or whether a therapeutic intervention would need to be additionally administered during the LRI or development of experimental asthma to ameliorate disease. Future studies should administer sgp130Fc protein or anti-IL-17A prophylactically and assess whether this would reverse disease susceptibility. Additionally, we did not identify the cellular source of IL-17A during the state of LGSI, when it was found to be elevated in the lung and liver. Intriguingly, the numbers of ILC3 were not elevated in the liver, suggesting that other lymphocytes, potentially γδ-T cells, are the likely source of IL-17A production. In summary, we demonstrated that a maternal HFD induces a state of LGSI in the infant and predisposes to sLRI and subsequent experimental asthma. Mechanistically, we demonstrated that the effects are mediated by an altered microbiome that drives the expansion of IL-6R-expressing neutrophils in the neonatal liver, and that the subsequent elevated sIL-6R expression, together with elevated IL-6 production, promotes disease pathogenesis via IL-6 trans-signaling.

## Methods

### Mouse strains and experimental design

Wild type C57BL/6J mice (from the Animal Resources Centre, WA, Australia), sgp130Fc mice (provided by Dr. Rose-John, University of Kiel), Ly6G^cre^ mice (Hasenberg, Hasenberg et al. 2015), IL-6Ra^fl/fl^ mice and IL-17A-deficient mice (both provided by Dr. Varelias, QIMR Berghofer MRI) (Nakae, Komiyama et al. 2002), and Ly6G^cre^IL-6Ra^fl/fl^ mice were housed and bred under specific pathogen-free conditions at the QIMR Berghofer Medical Research Institute (MRI) animal facility. Breeding age male and female mice were fed a control fat diet (CFD) or high fat diet (HFD) for 21 days prior to timed mating. The animals were maintained on the respective specialty diet for the duration of the experiment. The CFD and HFD chow was purchased from Specialty Feeds (WA, Australia). The CFD (Product code: SF13-081) and HFD (Product code: SF04-001) chow contained similar protein content (CFD = 23%; HFD = 22.6%), crude fiber (5.4%), and digestible energy (CFD = 15.6; HFD = 19 MJ/kg). However, the HFD had a higher fat content (23.5% by weight) in comparison to the CFD (5.3%) due to elevated amounts of lard, the CFD containing 23 g/kg and HFD containing 207 g/kg. All animal procedures were approved by the QIMR Berghofer MRI Animal Ethics Committee (approval number A1706-609M). Pneumonia virus of mice (PVM, strain J3666) was propagated as previously described (Sebina, Rashid et al. 2022). A single dose of 10 plaque forming unit (PFU) of PVM in 10 µl or the same volume of vehicle (10%, v/v, FCS in DMEM; Gibco) was administered via the intra-nasal (i.n.) route at postnatal day (PND)7 or PND10 (as indicated in the study design) (Loh, Simpson et al. 2020). To induce experimental asthma, mice were inoculated with PVM or vehicle at PND7, then exposed to low-dose cockroach allergen extract (CRE, Greer, USA; 1 µg i.n., 50 µl) or diluent (PBS) at PND 42, 49, 56 and 63 while under light isoflurane-induced anesthesia (Werder, Ullah et al. 2022). In the cross-fostering (CF) study, pups born to HFD-fed mothers were fostered by HFD- or CFD-fed mothers, and *vice-versa*. The fostered pups were removed from the biological mother and transferred to a different cage with the foster dam within 24 hours of birth. For the fecal microbiome transplant (FMT), a fecal suspension from CFD- or HFD-reared pups was oral gavaged to CFD- or HFD-reared pups at PND5, 7 and 9 as previously described (Sikder, Rashid et al. 2023). To deplete neutrophils, mice were treated intraperitoneally with anti-Ly6G antibody (25 mg/kg. bd. Wt; clone: 1A8; Bioxcell) or an isotype-matched control antibody at PND5, 7 and 9.

### Preparation of fecal suspension for the FMT

The entire gut was collected from culled CFD- and HFD-reared pups at PND7 and placed separately into 0.5 mL microcentrifuge tubes on ice. The tubes were immediately transferred to a Ruskin hypoxia chamber where the guts were longitudinally cut open and washed with sterile cold PBS and collected into a separate tube. The fecal suspension was vortexed for three mins at room temperature, passed through a 70 µm cell strainer followed by centrifugation at 800 g for two mins to remove any undissolved particulates. The supernatant was carefully collected and mixed with 50% (v/v) anaerobic glycerol in PBS and stored in serum bottles closed with a butyl rubber stopper at −80°C. Neonatal mice were later gavaged with the fecal suspension (10^8^ cfu in 50 μl) at PND5, 7, and 9. The following treatments and experimental groups were tested:

1. (F)H→H: FMT from HFD-reared pups to HFD-reared neonates.
2. (F)H→C: FMT from HFD-reared pups to CFD-reared neonates.
3. (F)C→H: FMT from CFD-reared pups to HFD-reared neonates.
4. (F)C→C: FMT from CFD-reared pups to CFD-reared neonates.

### Tissue collection

Following euthanasia, blood was obtained via cardiac puncture, spun at 13,000 rpm at 4°C for 10 mins, and serum collected and stored at −80°C. Bronchoalveolar lavage (BAL) fluid was collected as described previously (Werder, Ullah et al. 2022). Briefly, following euthanasia, the lungs of tracheotomized mice were washed with 400 µl of ice-cold PBS. The lavage fluid was then centrifuged at 13,000 rpm at 4°C for ten mins, and the cell-free supernatant immediately frozen on dry ice, prior to storage at −80°C. After removal of the thoracic region, the left lung and the post caval lobe were excised for immunophenotyping by flow cytometry. For histological analysis, the inferior right lung lobe was stored in 10% neutral buffered formalin overnight, then transferred to 70% ethanol for embedding in paraffin blocks. The superior and the middle right lobe was snap frozen in dry ice and stored at −80°C prior to proteomic analysis. Other excised organs/tissues, including the colon, small intestine, inguinal adipose tissue, and liver, were snap frozen and stored at −80°C prior to further analysis. For some experiments, the fecal content of the gut was collected, snap frozen, and stored at −80°C prior to DNA extraction.

### Cytokine quantification

Tissue samples stored in −80°C were thawed and weighed. The tissues were then homogenized with a tissue-tearor (BioSpec Products Inc) in protein assay buffer containing sodium chloride (150 mM), IGEPAL® CA-630 (1%), sodium deoxycholate (0.5%), sodium dodecyl sulphate (0.1%) and Tris buffer (50mM). The samples were then centrifuged at 13,000 rpm for 10 mins at 4°C, and the supernatant collected. The concentration of mouse IL-1β, IL-6, IL-17A, TSLP, IL-12p70, IL-23p19, IFN-γ, IL-25 (Biolegend, San Diego, CA), soluble IL-6R, IL-33, IFN-λ (R&D Systems, Minneapolis, MN), IL-4, and IL-5 (BD Biosciences) was measured by ELISA according to the manufacturer’s instructions. IL-13 concentration was measured by cytokine bead array (Becton Dickinson). For the purposes of genotyping, sgp130Fc was measured in serum using a custom ELISA system as previously described (Rabe, Chalaris et al. 2008). A list of ELISA reagents is provided in Supplementary Table S1.

### Flow cytometry

Lung lobes were mechanically dissociated into a single cell suspension by forcing the tissue through a 70 μm cell strainer in FACs buffer (2% fetal calf serum in PBS). The cell suspension was centrifuged then treated with Gey’s buffer to remove contaminating erythrocytes (Werder, Ullah et al. 2022). The cells were washed in FACs buffer, counted using a hemocytometer, then seeded into a v-bottom plate. Cells were washed in PBS and then stained for 10 minutes at room temperature with Zombie Aqua fixable viability dye (Biolegend, San Diego, CA) for dead cell exclusion. Cells were washed twice in PBS/2% FBS, incubated with anti-FcγRIII/II (Fc block) for 15 minutes at 4°C, washed again, and then stained using fluorescent-labeled antibodies directed against surface antigens for 30 minutes at 4°C. For intracellular staining, the cells were fixed and permeabilized using the BD Cytofix/Cytoperm™ kit as per the manufacturer’s instructions (BD Biosciences, San Jose, CA). Stained cells were washed twice and data were acquired on a BD LSR Fortessa X-20 (BD Biosciences, San Jose, CA), using FACS Diva Software v8 (BD Biosciences, San Jose, CA). Data were analyzed using FlowJo v10.6 (Tree Star Inc., Ashland, OR). To identify immune cells, gating was performed on live, single cells, and immuno-phenotyped as follows: Neutrophils: CD45^+^CD11b^+^Ly6G^+^; Eosinophils: CD45^+^Ly6G^-^CD11c^-^CD11b^+^SiglecF^+^; CD4^+^ Th1 cells: CD45^+^CD90.2^+^TCRβ^+^CD8^-^CD4^+^Tbet^+^; CD4^+^ Th2 cells: CD45^+^CD90.2^+^TCRβ^+^CD8^-^ CD4^+^ RORγt^-^GATA3^+^; CD4^+^ Th17 cells: CD45^+^CD90.2^+^TCRβ^+^CD8^-^CD4^+^GATA3^-^RORγt^+^; NK cells: CD45^+^CD90.2^+^TCRβ^-^NK1.1^+^Tbet^+^ NKp46^+^CD200R1^-^; ILC1: CD45^+^CD90.2^+^TCRβ^-^ NK1.1^+^Tbet^+^NKp46^+^CD200R1^+^ (Weizman, Adams et al. 2017); ILC2: CD45^+^CD90.2^+^TCRβ^-^ NK1.1^-^RORγt^+^GATA3^+^; ILC3: CD45^+^CD90.2^+^TCRβ^-^NK1.1^-^ GATA3^+^RORγt^+^; Classical monocytes: CD45^+^CD11b^+^CD11c^-^CD115^+^Ly6C^+^; conventional dendritic cell type-1 (cDC1): MHCII^+^CD11c^+^CD26^+^CD64^-^CD172^-^XCR1^+^; cDC2: MHCII^+^CD11c^+^CD26^+^CD64^-^ CD172^+^XCR1^-^MAR1^-^; Monocyte derived DC (moDC): MHCII^+^ CD11c^+^CD26^-^CD64^+^XCR1^-^ CD172^+^MAR1^+^. The list of antibodies used for flow cytometry is summarized in Supplementary Table S2.

### Histology and Immunohistochemistry

Paraffin embedded lung lobes were sectioned and immunohistochemistry (IHC) for PVM, MUC5ac, and α-smooth muscle actin performed as described previously (Werder, Lynch et al. 2018, Curren, Ahmed et al. 2023). The list of antibodies used is summarized in Supplementary Table S3. Sections were scanned using Scanscope XT software and the mucus score, airway smooth muscle area, and number of PVM-immunoreactive AECs quantified as previously described (Sikder, Rashid et al. 2023).

### Primary hepatocyte and neutrophil isolation and co-culture studies

Hepatocytes were isolated from the liver of neonatal mice reared by CFD- and HFD-fed mothers as based on modified enzymatic digestion and percoll density separation as described previously (Korelova, Jirouskova et al. 2019). In brief, following euthanasia, the heart was perfused with cold PBS to remove erythrocytes from the liver. The liver was then excised, placed in Roswell Park Memorial Institute (RPMI) 1640 media supplemented with Liberase^TM^, and incubated at 37°C for 20 mins with continuous shaking. After the digestion process, the liver was cut into small pieces using a sterile scalpel and filtered through a 70 μm cell strainer. The cell suspension was centrifuged at 50 g for 5 mins followed by red blood cell lysis using Gey’s buffer for 10 minutes. The samples were then centrifuged at 50 g for 5 minutes and the pellet washed with Hank’s balanced salt solution (HBSS). A gradient separation step using 40% percoll in HBSS was performed by centrifugation at 100 g for 10 min with no brake. The cell pellet was resuspended in DMEM (high glucose) containing 10% FCS and 2% Penicillin/Streptomycin (200 U and 200 µg/ml, respectively), and the cells seeded into a collagen precoated 96 well plate at 200,000 cells/well. 24 hrs later the plate was centrifuged and supernatant harvested and stored at −80°C until use. To isolate the neutrophils from the liver, the tissue was digested as described above, and the single cell suspension spun at 50 g to sediment the hepatocytes. The cells in suspension were collected and layered over 33% Percoll in PBS. Following centrifugation at 600 g for 15 mins with the brake off, the neutrophil-rich pellet was collected, and the contaminating red blood cells lysed by incubation in Gey’s lysis buffer for 2 mins, before further wash steps and in 2% FCS in PBS. The cell pellet was incubated with anti-FcγRIII/II (Fc block) for 15 minutes at 4°C, then stained with a cocktail of antibodies to allow for identification (CD3^-^CD19^-^NK1.1^-^CD45^+^CD11c^-^ CD11b^+^SiglecF^-^Ly6G^+^) and sorting on a BD FACSAria^TN^ III cell sorter. The neutrophils (>98% purity) were then cultured in the absence or presence of the hepatocyte supernatant. After 48 hr, the plate was centrifuged, and the neutrophil supernatant harvested for quantification of sIL-6R by ELISA.

### Profiling of fecal microbiomes

DNA was extracted from 200 mg of each fecal sample via bead beating and column purification using a Maxwell 16 MDx Instrument (Promega, USA), as previously described (Sikder, Rashid et al. 2023). Universal 16S rRNA genes were then amplified by polymerase chain reaction using the primers 926F 5’-AAA CTY AAA KGA ATT GRC GG-3’ and 1392wR 5’-ACG GGC GGT GWG TRC-3’ (Engelbrektson, Kunin et al. 2010) and sequenced using an Illumina MiSeq as previously described (Sikder, Rashid et al. 2023). Negative kit and PCR controls were included to confirm the absence of contamination. Sequence data were then grouped into operational taxonomic units (OTUs) sharing 97% similarity, assigned Silva 138 taxonomy (Quast, Pruesse et al. 2012) and summarised in an OTU table using a modified UPARSE pipeline as described previously (Sikder, Rashid et al. 2023). The OTU table was then rarefied to 1150 reads per sample, Hellinger transformed, and subjected to permutational multivariate analysis of variance (PERMANOVA) to determine whether the composition of fecal bacterial communities differed significantly between levels of the factor maternal diet (CFD, HFD). OTUs that were significantly associated with diet were identified using Dufrene-Legendre Indicator Species Analysis as implemented by the indval function within the R package labdsv (Dufrêne and Legendre 1997).

### Statistical analysis

All data were represented as the mean ± standard error of the mean (SEM) or box and whiskers plots; the box represents the quartiles, and the whiskers represent the range. The data were analyzed by using either a Mann-Whitney *U* test (for two groups) or a One- or Two-way ANOVA (for multiple groups) with Tukey’s post-hoc test. Statistical analyses were performed using GraphPad Prism (Version 8; GraphPad Software, La Jolla, California). The figure legends contain information as to the specific statistical tests used. Significance levels were determined as follows: * indicates p < 0.05, ** indicates p < 0.01, and *** indicates p < 0.001.

## Supporting information

Graphical Summary

Publication License

## Author Contribution

S.P. conceived the project and designed the experiments, which were primarily performed by B.C. and T.A. with support from R.B.R., I.S., M.A.A.S, D.H., M.A., M.A.U., A.B., M.M.R. Data analysis and presentation was performed by B.C. and T.A. Critical analyses and tools were provided by A.V., M.A.P., G.A.R., S.R.-J., R.H., P.Ó C, K.M.S. and P.G.D. The first draft of the manuscript was written by S.P. with support from T.A. All the authors reviewed, edited, and approved the final manuscript.

## Acknowledgement

This work was supported by a National Health and Medical Research Council of Australia grant awarded to S.P., P.Ó C, and P.G.D (1141581). The work of S.R.-J. was supported by the Deutsche Forschungsgemeinschaft Bonn, Germany in the Cluster of Excellence “Precision Medicine in Chronic Inflammation” (grant no. EXC2167). The authors thank all the staff from the QIMR Berghofer animal facility, flow cytometry laboratory, and histology facility for their assistance.

## Abbreviations

AEC: Airway epithelial cell
ASM: Airway smooth muscle
BAL: Broncho-alveolar lavage
CFD: Control fat diet
CRE: Cockroach allergen extract
FACs: Fluorescence-Activated Cell Sorting
FMT: Fecal microbiome transplants
HFD: High fat diet
IHC: Immunohistochemistry
ILC1: Group 1 innate lymphocyte cells
ILC2: Group 2 innate lymphocyte cells
ILC3: Group 3 innate lymphocyte cells
LGSI: Low grade systemic inflammation
NK: Natural killer cells
PND: Postnatal day
PVM: Pneumonia virus of mice
RSV: Respiratory syncytial virus
sIL-6R: Soluble IL-6 receptor
sLRI: Severe lower respiratory infection
sgp130: Soluble gp130

---

**Figure S1.**
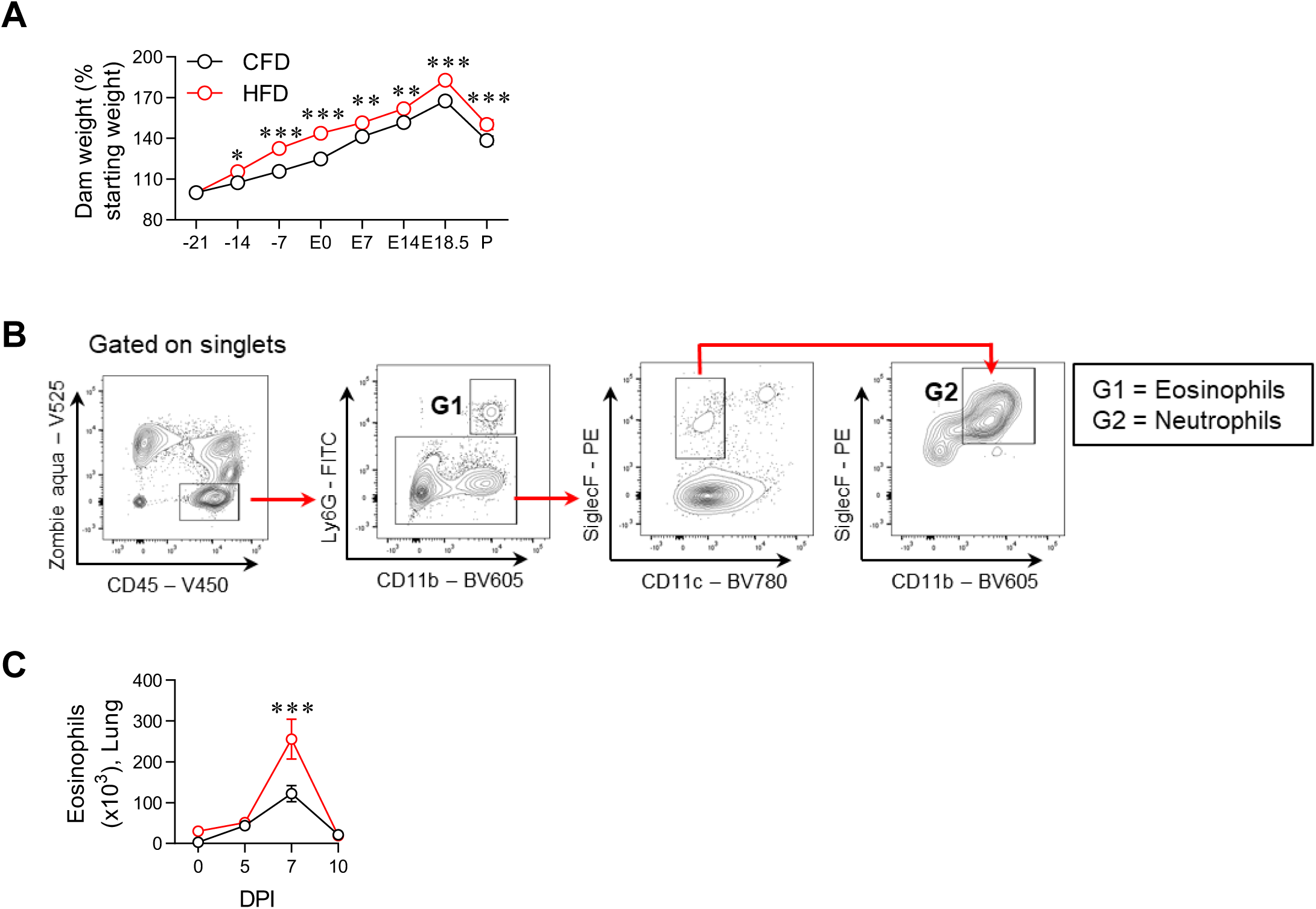
Weight gain of CFD- and HFD-fed dams, gating strategy to identify granulocytes and eosinophils in PVM infected neonates. **(A)** Maternal bodyweight (as percentage of starting weight) prior to timed-mating (−21 to E0), during gestation (E0-E18.5) and at partum (P). **(B)** Gating strategy to identify neutrophils and eosinophils. **c** Eosinophils in the lung. Data are shown as mean ± SEM. Data are pooled from 2 independent experiments, *n* = 3-4 mice per group in each experiment. *P* values (*p< 0.05, **p< 0.01, ***p< 0.001) were derived by two-way ANOVA with Tukey’s post hoc test.

**Figure S2.**
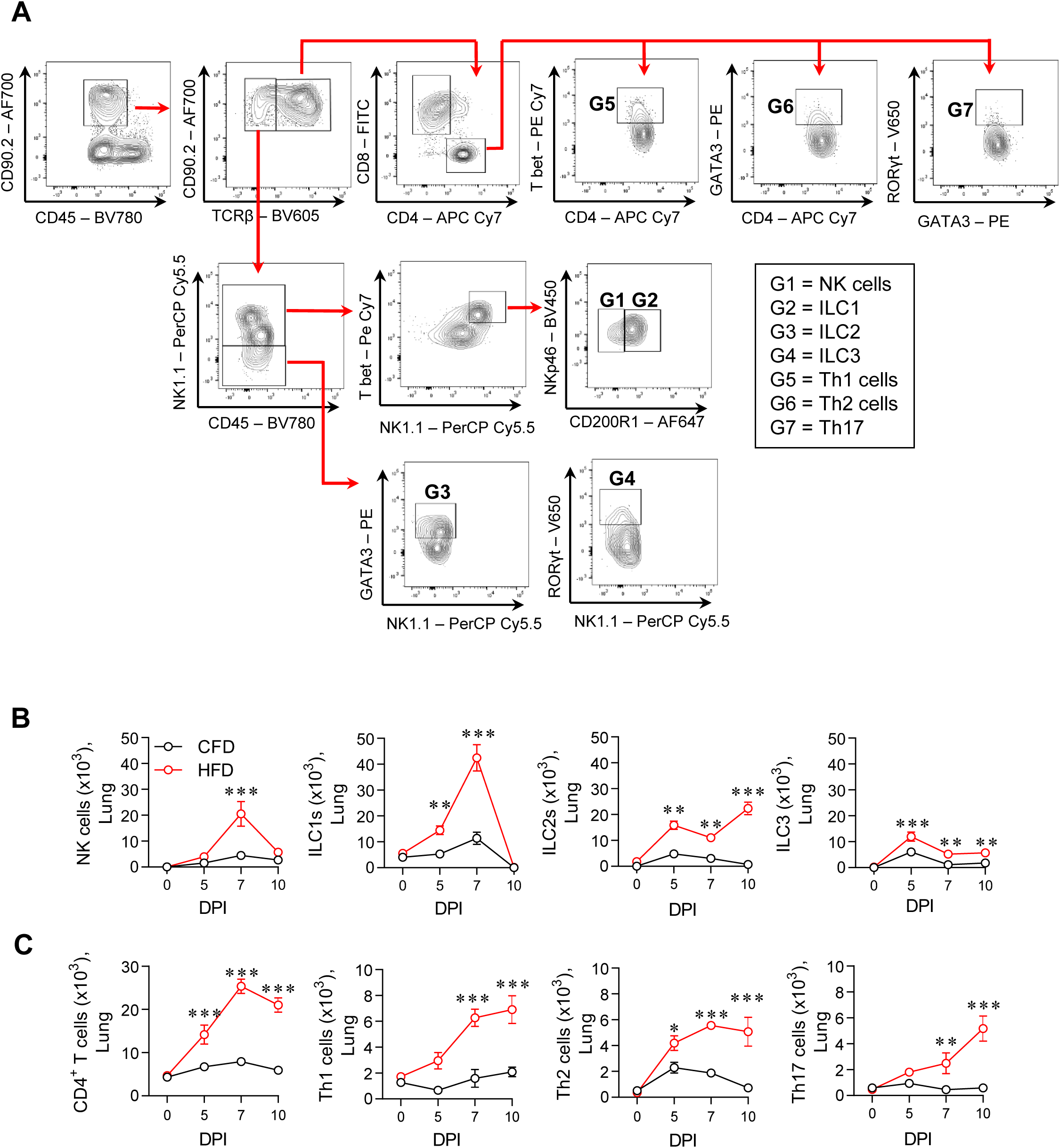
Gating strategy to identify CD4^+^ T helper cells, innate lymphoid cells (ILC) and their numbers in the lung of CFD- and HFD-reared neonates following PVM infection. **(A)** Gating strategy to identify CD4^+^ T helper cell and ILC subsets. **(B)** Natural killer (NK) cells (first from left), group-1 innate lymphoid cells (ILC1s), ILC2s and ILC3s in the lung post PVM infection. (**C)** CD4^+^ T helper cells, CD4^+^ T helper type-1 (Th1), Th2 and Th17 cells in the lung post PVM infection. Data are shown as mean ± SEM. Data are pooled from 2 independent experiments, *n* = 3-4 mice per group in each experiment. *P* values (*p< 0.05, **p< 0.01, ***p< 0.001) were derived by two-way ANOVA with Tukey’s post hoc test.

**Figure S3.**
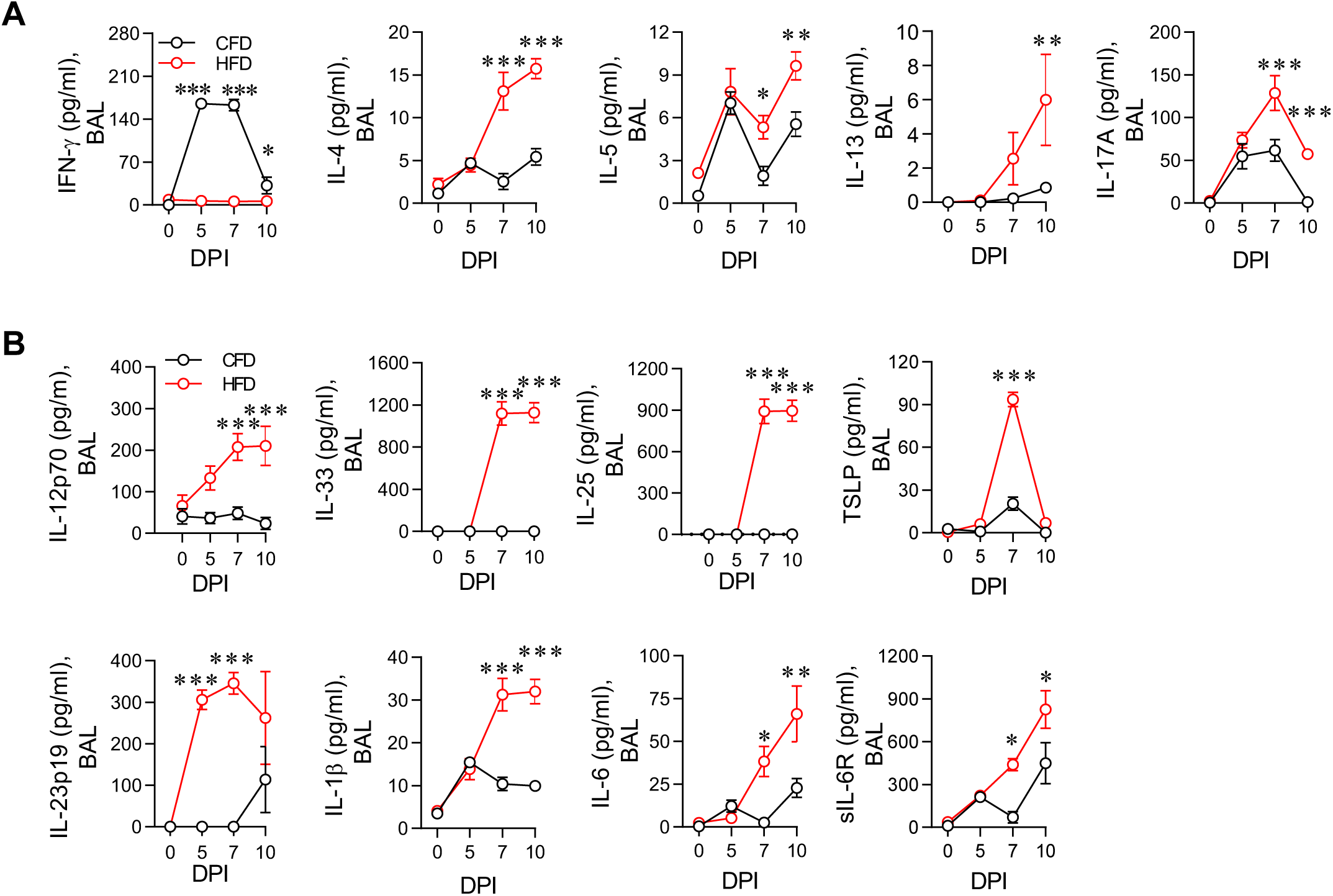
Elevated instructive and effector cytokines in the BAL fluid of HFD-reared neonates following PVM infection. **(A)** IFN-γ, IL-4, IL-5, IL-13 and IL-17A. **(B)** IL-12p70, IL-33, IL-25, Thymic Stromal Lymphopoietin (TSLP) IL-23p19, IL-1β, IL-6 and sIL-6R in BAL fluid. Data are shown as mean ± SEM. Data are pooled from 2 independent experiments, *n* = 3-4 mice per group in each experiment. *P* values (*p< 0.05, **p< 0.01, ***p< 0.001) were derived by two-way ANOVA with Tukey’s post hoc test.

**Figure S4.**
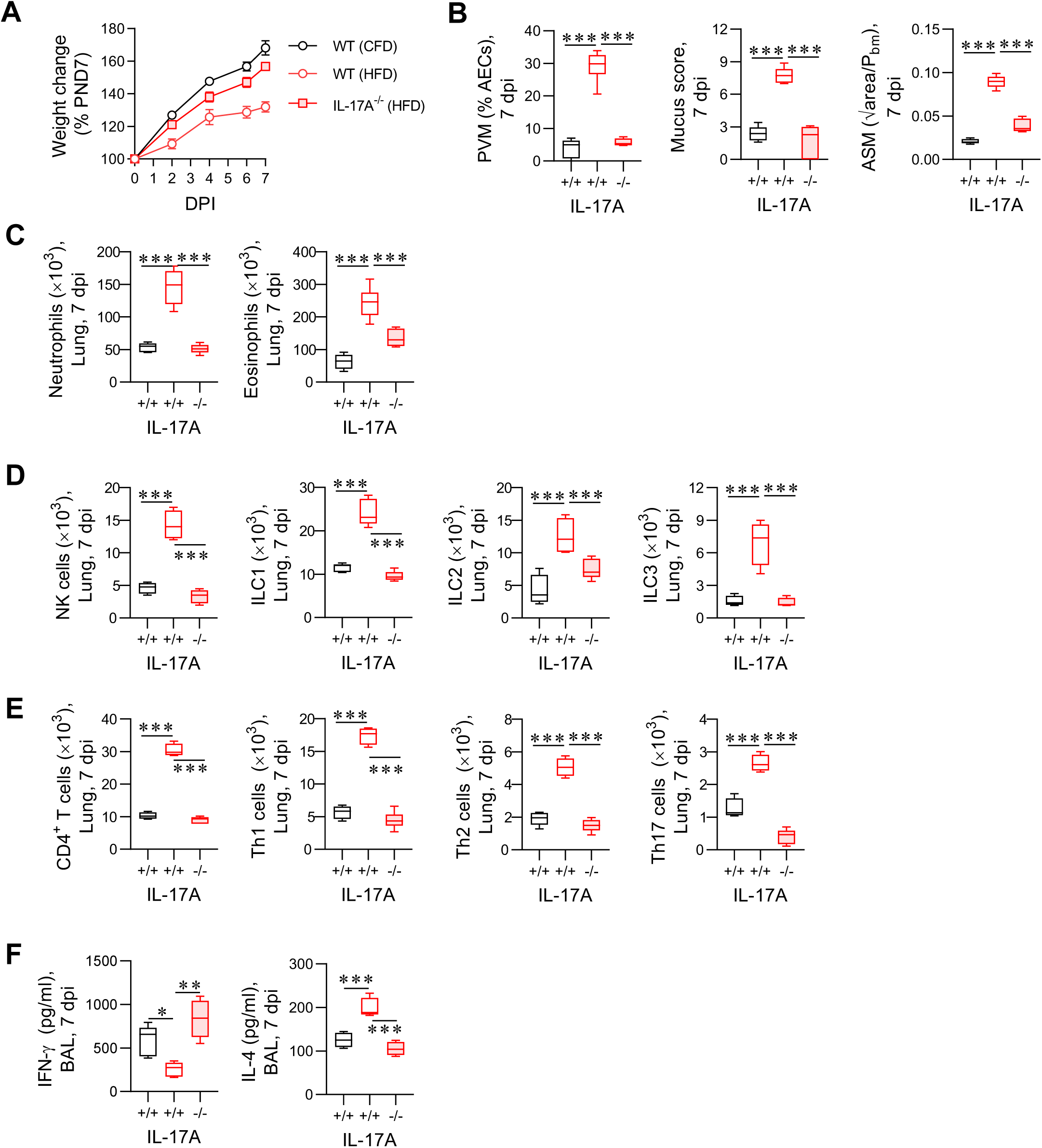
HFD-reared IL-17A-depleted neonates develop a less severe LRI following challenge with PVM. **(A)** Weight change in pups post PVM inoculation, expressed as percentage of weight at postnatal day (PND7). **(B)** Viral load, Mucus score, and ASM area. **(C)** Neutrophils (left panel) and eosinophils in the lung. **(D)** Natural killer (NK) cells, ILC1s, ILC2s and ILC3s. **(E)** Total CD4^+^ T helper cells, CD4^+^ Th1, Th2 and Th17 cells in the lung. **(F)** IFN-γ, IL-4 and IL-17A in BAL fluid. Data are shown as mean ± SEM or box-and-whisker plots (median, quartiles, and range). Data are pooled from 2 independent experiments, *n* = 3-4 mice per group in each experiment. *P* values (*p< 0.05, **p< 0.01, ***p< 0.001) were derived by one-way or two-way ANOVA with Tukey’s post hoc test.

**Figure S5.**
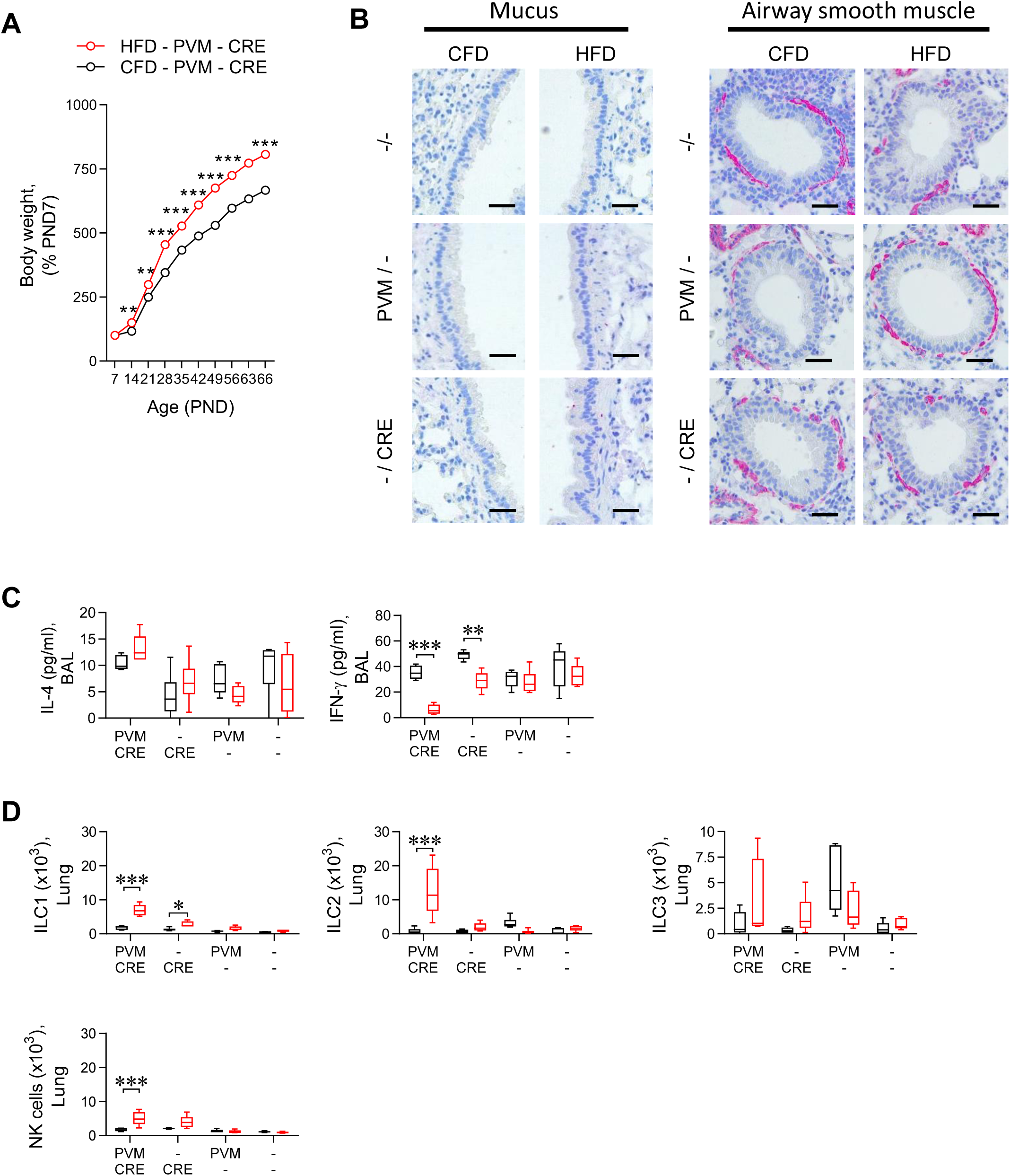
A maternal high fat diet (HFD) and severe bronchiolitis in the progeny predisposes to subsequent asthma in later life. **(A)** Weight of mice as percentage of PND7. **(B)** Representative histology images of secreting protein muc5ac (pink) in the airway (left panel) and ASM surrounding the bronchioles (pink), ×20 magnification, scale bar = 100 µm. **(C)** IL-4 and IFN-γ in BAL fluid. **(D)** ILC1s, ILC2s, ILC3s and NK cells. Data are shown as mean ± SEM or box-and-whisker plots (median, quartiles, and range). Data are pooled from 2 independent experiments, *n* = 3-4 mice per group in each experiment. *P* values (*p< 0.05, **p< 0.01, ***p< 0.001) were derived by two-way ANOVA with Tukey’s post hoc test.

**Figure S6.**
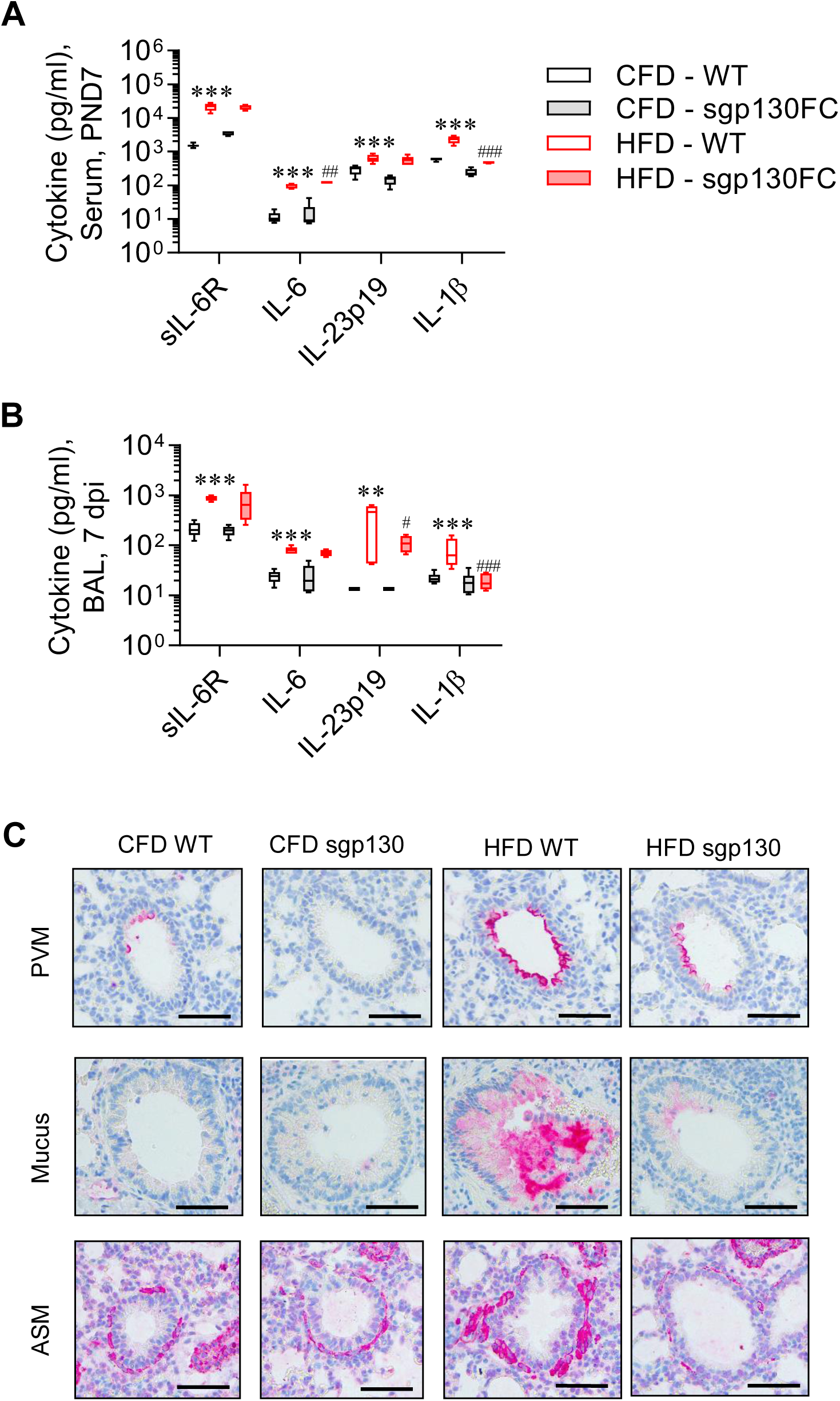
HFD-reared sgp130Fc transgene mice are protected from LGSI, sLRI, and subsequent asthma. **(A-B)** sIL-6R, IL-6, IL-23p19 and IL-1β in (A) serum at PND7 and (B) BAL fluid at 7 dpi. **(C)** Representative histology images of viral load (top panel), muc5ac protein and airway smooth muscle (ASM) (bottom panel), ×20 magnification, scale bar = 100 µm. Data are shown as box-and-whisker plots (median, quartiles, and range). Data are pooled from 2 independent experiments, *n* = 3-4 mice per group in each experiment. *P* values (*p< 0.05, **p< 0.01, ***p< 0.001) were derived by one-way ANOVA with Tukey’s post hoc test. * were used to compare CFD_WT vs HFD_WT, # used to compare HFD_WT vs HFD_sgp130Fc Tg mice.

**Figure S7.**
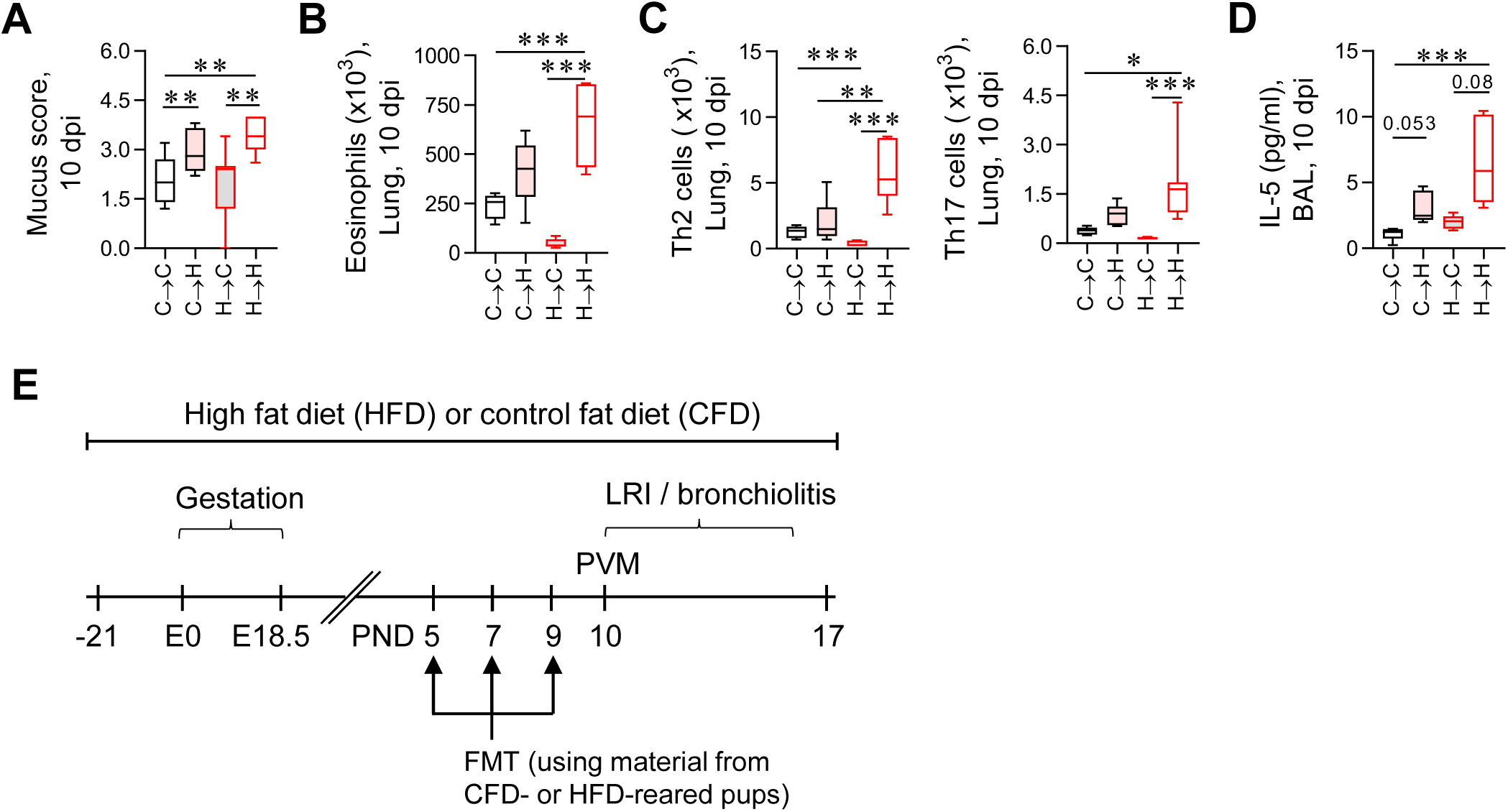
Effect of cross fostering on the immune response during PVM infection. **(A-D)** Cross-fostering groups: pups moved from a CFD-fed dam to a HFD-fed dam (C→H), pups moved from a HFD-fed dam to a HFD-fed dam (H→H), pups moved from a CFD-fed dam to a CFD-fed dam (C→C) and pups moved from a HFD-fed dam to a CFD-fed dam (H→C). (A) Mucus score. (B) Eosinophils in the lung. (C) CD4^+^ Th2 and Th17 cells. (D) IL-5 in BAL fluid at 10 dpi. **e** Study design of fecal microbiota transplant (FMT) experiment. Data are shown as box-and-whisker plots (median, quartiles, and range). Data are pooled from 2 independent experiments, *n* = 3-4 mice per group in each experiment. *P* values (*p< 0.05, **p< 0.01, ***p< 0.001) were derived by one-way or two-way ANOVA with Tukey’s post hoc test.

**Figure S8.**
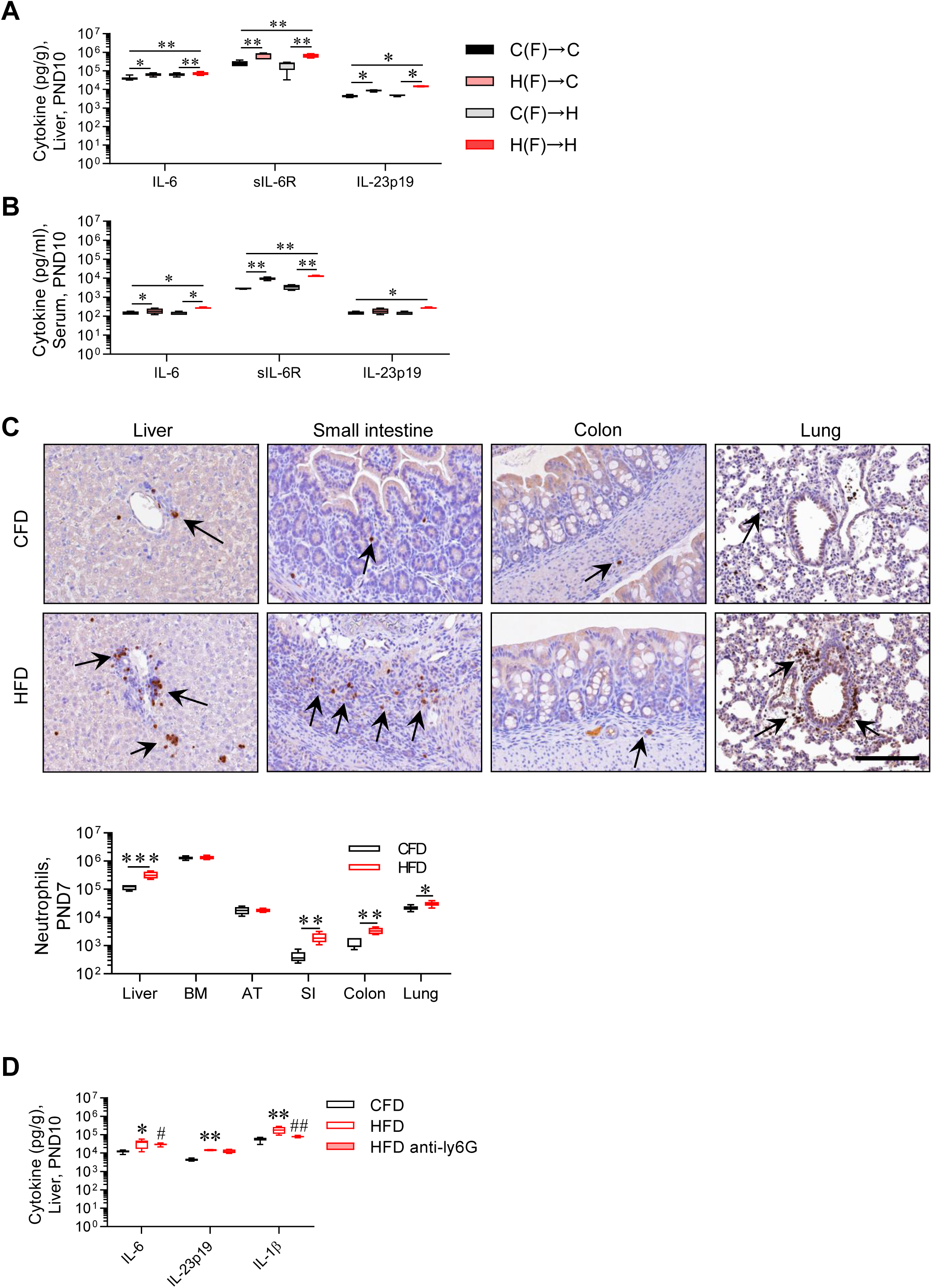
Attenuated type-17 instructive cytokine in HFD-reared neonates after FMT from CFD pups and neutrophil depletion. **(A-B)** IL-6, sIL-6R, IL-23p19 expression in the (A) liver and (B) serum after FMT PND10. **(C)** Representative histology images of neutrophils (brown, indicated by black arrow) in the liver, SI, colon and lung (Top panel), ×20 magnification, scale bar = 100 µm and neutrophil numbers in HFD- and CFD-reared neonatal liver, bone marrow (BM), adipose tissue (AT), small intestine (SI), colon and lung at PND7. **(D)** IL-6, IL-23p19 and IL-1β expression in the liver after neutrophil depletion at PND10. Data are shown as box-and-whisker plots (median, quartiles, and range). Data are pooled from 2 independent experiments, *n* = 3-4 mice per group in each experiment. *P* values (*p< 0.05, **p< 0.01, ***p< 0.001) were derived by one-way or two-way ANOVA with Tukey’s post hoc test.

**Figure S9.**
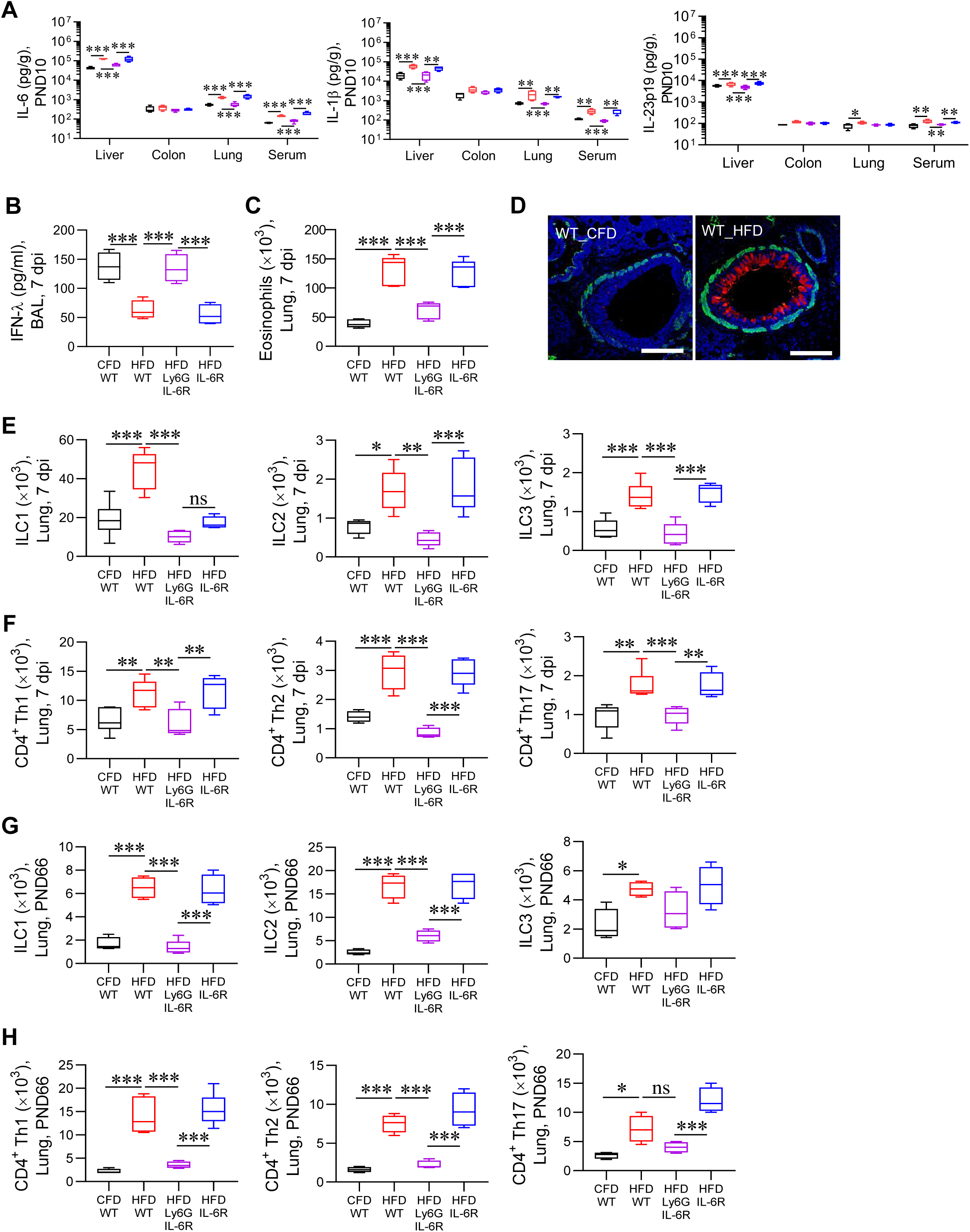
Ly6G^cre^IL-6Ra^fl/fl^ neonates are protected from LGSI, sLRI, and the subsequent development of asthma. **(A)** IL-6, IL-1β and IL-23p19 levels in the liver, colon, lung and serum at PND10. **(B)** IFN-λ in BAL fluid. **(C)** Eosinophils in the lung. **(D)** Representative immunofluorescence image of Muc5ac protein (Red) and airway smooth muscle (ASM; Green), ×20 magnification, scale bar = 100 µm. **e** ILC1s, ILC2s, ILC3s. **f** CD4^+^ Th1, Th2 and Th17 cells at 7 dpi. **(G-H)** Asthma phase. (G) ILC1, ILC2, ILC3 and (H) Th1, Th2, Th17 cells in the lung at PND66. Data are shown as box-and-whisker plots (median, quartiles, and range). Data are pooled from 2 independent experiments, *n* = 3-4 mice per group in each experiment. *P* values (*p< 0.05, **p< 0.01, ***p< 0.001) were derived by one-way or two-way ANOVA with Tukey’s post hoc test.

**Table S1:** Reagents for ELISA.

| **Name** | **Catalogue No.** | **Source** |
| --- | --- | --- |
| IFN-γ | 430815 | Biolegend |
| IFN-λ | RDSDY1789B | R&D Systems |
| sIL-6Ra | RDSDY1830 | R&D Systems |
| IL-4 | Capture: 554434<br>Detection: 554390 | BD Biosciences |
| IL-5 | Capture: 554393<br>Detection: 554397 | BD Biosciences |
| IL-13 | 562273 | BD Biosciences |
| IL-6 | 431301 | Biolegend |
| IL-12p70 | 433604 | Biolegend |
| IL-23p19 | 431601 | Biolegend |
| IL-17a | 432501 | Biolegend |
| IL-25 | RDSDY1399 | R&D Systems |
| IL-33 | RDSDY3626 | R&D Systems |
| TSLP | 434104 | Biolegend |
| IL-1β | 432601 | Biolegend |

**Table S2:** Fluorochrome conjugated antibodies for flow cytometry.

| **Name** | **Catalogue No.** | **Clone** | **Source** |
| --- | --- | --- | --- |
| Ly6G-FITC | 551460 | IA8 | BD Bioscience |
| NK1.1-PerCpCy5.5 | 156526 | PK136 | Biolegend |
| CD11b-BV605 | 101257 | M1/70 | Biolegend |
| CD11c-BV785 | 117336 | N418 | Biolegend |
| Siglec-F-PE | 552126 | E50-2440 | BD Bioscience |
| Zombie aqua Fixable Viability Kit | 423102 |  | Biolegend |
| Ly6C-BV570 | 128030 | HK1.4 | Biolegend |
| CD115-BV605 | 135517 | AFS98 | Biolegend |
| CD8-FITC | 100706 | 53-6.7 | Biolegend |
| CD200R1-APC | 566345 | OX-110 | BD Bioscience |
| CD90.2-AF700 | 140324 | 53-2.1 | Biolegend |
| CD4-APCCy7 | 552051 | GK1.5 | BD Bioscience |
| Nkp46-BV421 | 137612 | 29A1.4 | Biolegend |
| TCRβ-BV605 | 109241 | H57-597 | Biolegend |
| RORγt-BV650 | 564722 | Q31-378 | BD Horizon |
| CD45-BV421/BV785 | 103133/103149 | 30-F11 | Biolegend |
| GATA3-PE | 12-9966-42 | TWAJ | Invitrogen |
| Tbet-PECy7 | 644824 | 4B10 | Biolegend |
| MHCII- APCCy7 | 107628 | M5/114.15.2 | Biolegend |
| CD64-BV421 | 139309 | X54-5/7.1 | Biolegend |
| CD26-PE/Cyanine7 | 137810 | DPP-4 | Biolegend |
| CD172a-AF700 | 144022 | P84 | Biolegend |
| XCR1-BV650 | 148220 | ZET | Biolegend |
| FceR1 alpha<br>MAR1-PE | 12-5898-82 | MAR-1 | eBioscience |

**Table S3:** Antibodies used for Immunohistochemistry and Immunofluorescence imaging:

| **Antibodies** | **Catalogue No.** | **Source** |
| --- | --- | --- |
| Anti-α-smooth muscle actin antibody, clone 1A4 | A5228-100UL | Sigma |
| Anti-Muc5ac antibody | MA512178 | ThermoFisher |
| Primary antibody for PVM | A kind gift from by Dr. Ursula Buchholz, Laboratory of Infectious Diseases, National Institute of Allergy and Infectious Diseases |  |
| Anti-mouse Ly6G (abcam) |  | In house |
| Anti-Rabbit IgG (whole molecule)–Alkaline Phosphatase | A9919-.5ML | Sigma |
| Anti-Mouse IgG (whole molecule)-Alkaline Phosphatase | A3562-.5ML | Sigma |
| Anti-Mouse AF555 (Secondary Ab) | A-21422 | Life technologies Australia Pty. Ltd. |
| Anti-Mouse AF488 (Secondary Ab) |  | In house |

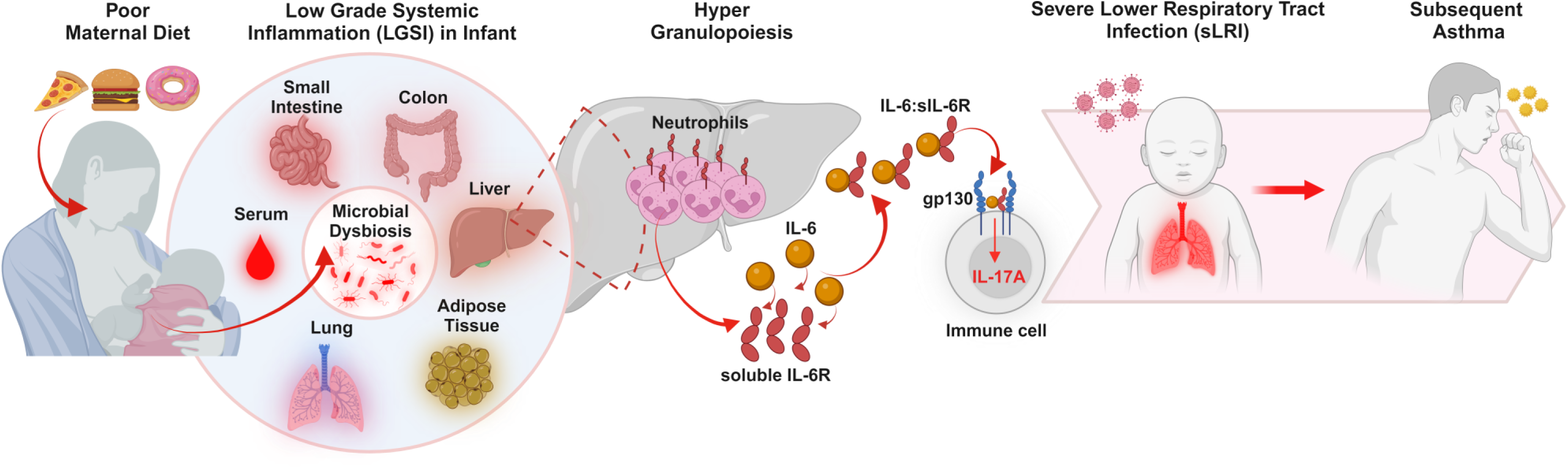

