## Supplementary figures and images for "A maternal high-fat diet predisposes to infant lung disease via increased neutrophil-mediated IL-6 trans-signaling"

### Graphical Summary

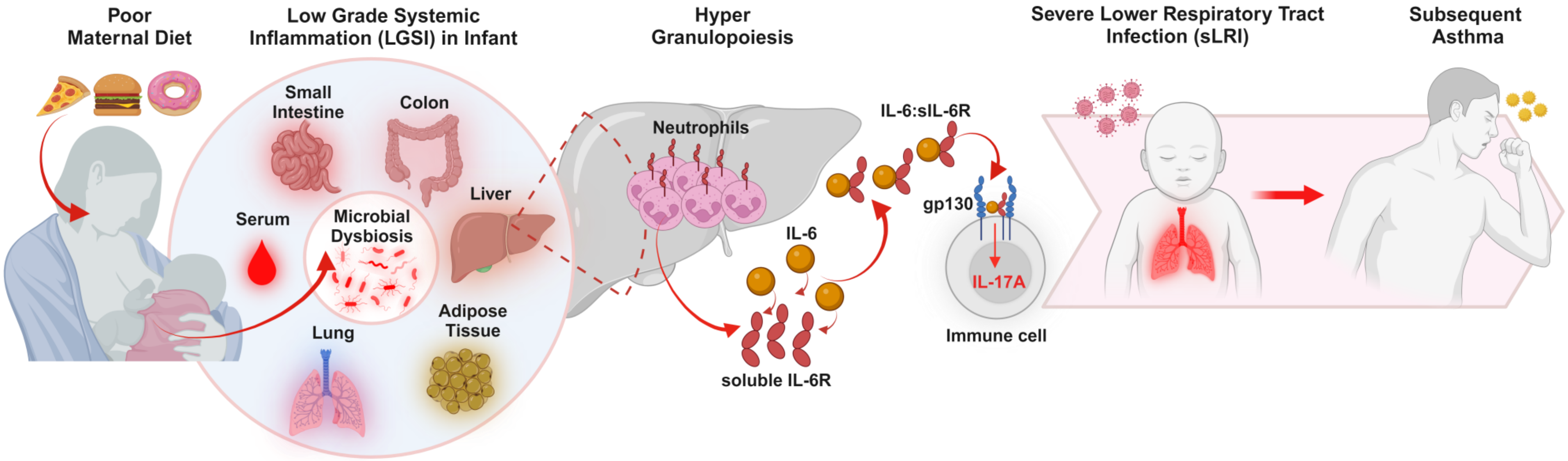
